# Mitogenomic Phylogeny of *Callithrix* with Special Focus on Human Transferred Taxa

**DOI:** 10.1101/2020.08.12.247692

**Authors:** Joanna Malukiewicz, Reed A. Cartwright, Nelson H.A. Curi, Jorge A. Dergam, Claudia S. Igayara, Silvia B. Moreira, Camila V. Molina, Patricia A. Nicola, Angela Noll, Marcello Passamani, Luiz C.M. Pereira, Alcides Pissinatti, Carlos R. Ruiz-Miranda, Daniel L. Silva, Anne C. Stone, Dietmar Zinner, Christian Roos

## Abstract

*Callithrix* marmosets are a relatively young primate radiation, whose phylogeny is not yet fully resolved. These primates are naturally para- and allopatric, but three species with highly invasive potential have been introduced into the southeastern Brazilian Atlantic Forest by the pet trade. There, these species hybridize with each other and endangered, native congeners. We aimed here to reconstruct a robust *Callithrix* phylogeny and divergence time estimates, and identify the biogeographic origins of autochthonous and allochthonous *Callithrix* mitogenome lineages. We sequenced 49 mitogenomes from four species (*C. aurita, C. geoffroyi, C. jacchus, C. penicillata*) and anthropogenic hybrids (*C. aurita* x *Callithrix* sp., *C. penicillata* x *C. jacchus, Callithrix* sp. x *Callithrix* sp., *C. penicillata* x *C. geoffroyi*) via Sanger and whole genome sequencing. We combined these data with previously published *Callithrix* mitogenomes to analyze five *Callithrix* species in total.

**Results:** We report the complete sequence and organization of the *C. aurita* mitogenome. Phylogenetic analyses showed that *C. aurita* was the first to diverge within *Callithrix* 3.54 million years ago (Ma), while *C. jacchus* and *C. penicillata* lineages diverged most recently 0.5 Ma as sister clades. MtDNA clades of *C. aurita, C. geoffroyi*, and *C. penicillata* show intraspecific geographic structure, but *C. penicillata* clades appear polyphyletic. Hybrids, which were identified by phenotype, possessed mainly *C. penicillata* or *C. jacchus* mtDNA haplotypes. The biogeographic origins of mtDNA haplotypes from hybrid and allochthonous *Callithrix* were broadly distributed across natural *Callithrix* ranges. Our phylogenetic results also evidence introgression of *C. jacchus* mtDNA into *C. aurita*.

**Conclusion:** Our robust *Callithrix* mitogenome phylogeny shows *C. aurita* lineages as basal and *C. jacchus* lineages among the most recent within *Callithrix*. We provide the first evidence that parental mtDNA lineages of anthropogenic hybrid and allochthonous marmosets are broadly distributed inside and outside of the Atlantic Forest. We also show evidence of cryptic hybridization between allochthonous *Callithrix* and autochthonous *C. aurita*. Our results encouragingly show that further development of genomic resources will allow to more clearly elucidate *Callithrix* evolutionary relationships and understand the dynamics of *Callithrix* anthropogenic introductions into the Brazilian Atlantic Forest.

## Background

*Callithrix* species represent a relatively young radiation, and divergence among lineages within the genus is estimated to be between approximately 0.7 and 2.5 million years ago (Ma) [1, 2,3]. Two major subgroups occur within the genus, the *aurita* group (*C. aurita*/*C. flaviceps*) and the *jacchus* group (*C. geoffroyi*/*C. kuhlii*/*C. jacchus*/*C. penicillata*), but the phylogeny of these lineages is not yet fully resolved. *Callithrix* species are naturally para- and allopatric across the Brazilian Atlantic Forest, Cerrado, and Caatinga biomes (Figure 1) [4,5], and natural hybridization occurs between some species [6]. However, *C. geoffroyi, C. jacchus*, and *C. penicillata* have high invasive potential [7,8] and have spread widely outside of their native ranges due to the legal and illegal pet trades. These species have established several allochthonous populations in the southeastern Brazilian Atlantic Forest [6,9,10] and hybridize with other allochthonous and autochthonous congeners [6,9,11,10], including endangered *C. aurita* and *C. flaviceps* [12,13]. Yet, determining evolutionary relationships between autochthonous, allochthonous, and hybrid *Callithrix* populations across Brazil is complicated by the unresolved *Callithrix* phylogeny.

**Figure 1.**
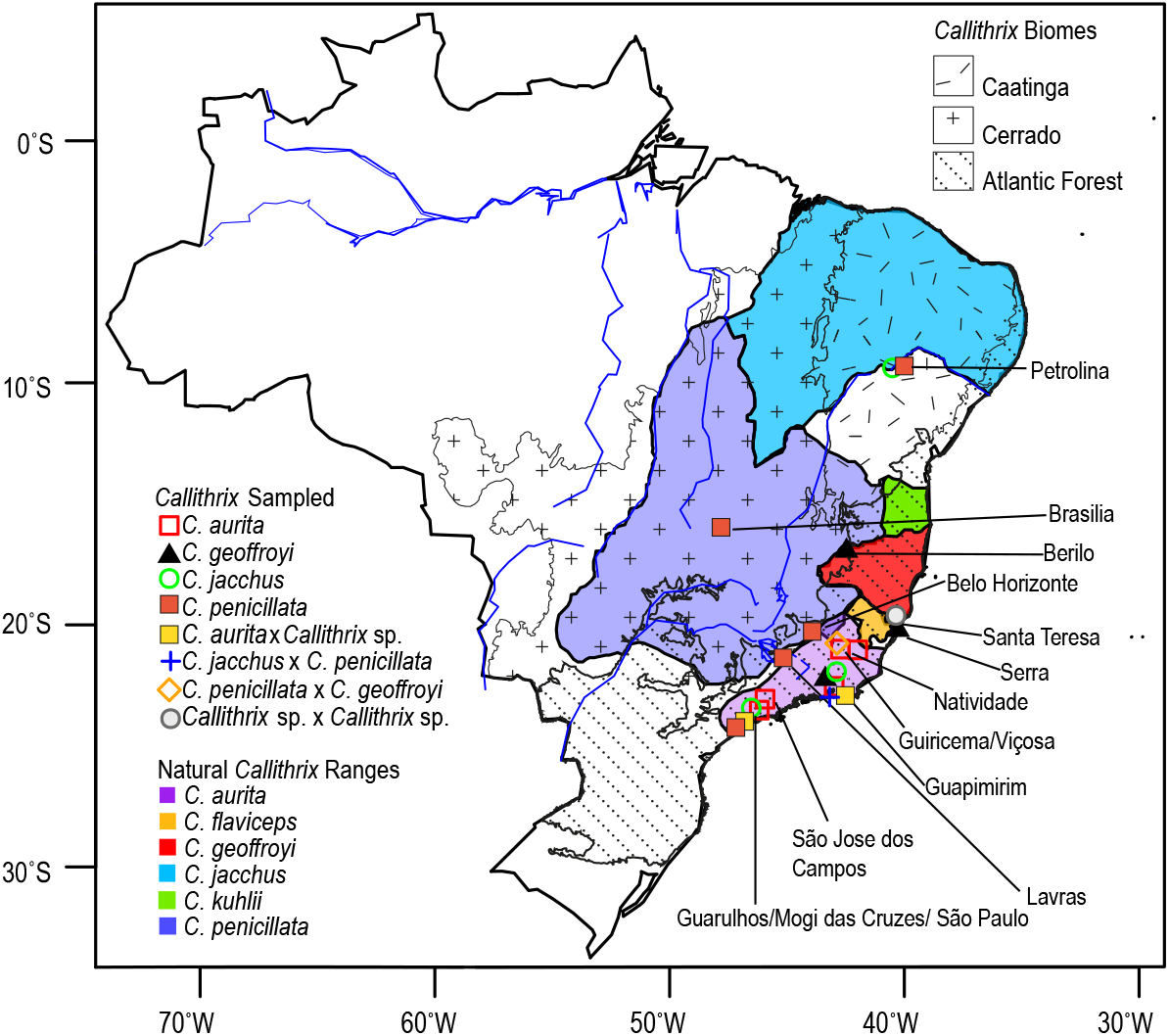
Approximate distribution of *Callithrix* species in Brazil (2012 IUCN Red List Spatial Data; http://www.iucnredlist.org/technical-documents/spatial-data) and geographic origins of Brazilian samples, as indicated by capital letter symbols. Locations of three biomes where *Callithrix* occur naturally, the Caatinga, Cerrado, and Atlantic Forest, are also indicated.

In general, mitochondrial DNA (mtDNA) can be utilized for an initial look into evolutionary relationships among taxa [e.g., 14,15] as well as track dispersal and gene flow patterns of allochthonous species [16]. MtDNA sequence data can also provide initial genetic insight in the direction of introgression (if sex-biased) when two species hybridize due to incongruences between phenotypes and haplotypes [e.g., 14,15]. The effective population size of mtDNA is one quarter of that of nuclear DNA from a diploid, bisexual population, which allows mtDNA lineages to coalescence relatively more quickly [17]. MtDNA is also considered a relatively fast mutating genetic marker [18]. As a result, lineage sorting and reciprocal monophyly are expected to occur faster in mtDNA than nuclear DNA, which can provide insight into shallow evolutionary relationships expected for young radiations.

One major challenge in applying genetic and genomic methods in *Callithrix* studies is an overall lack of genomic resources and sample material for most *Callithrix* species. Studies of *Callithrix* species have utilized mtDNA markers that generally resulted in polytomies and/ or poorly supported branching patterns, as well as polyphyly for *C. penicillata* and *C. kuhlii* [19,20,21,22,23]. Also, the few available genetic studies of allochthonous and hybrid *Callithrix* within the Atlantic Forest, all conducted within Rio de Janeiro state, used portions of mtDNA or the Y-chromosome that could not fully resolve the evolutionary relationships of *Callithrix* lineages [e.g., 11,23]. Nonetheless, [24] obtained a well-resolved phylogeny for the *jacchus* group using complete mitogenomes, but they only sampled one individual/species with unknown provenances.

To build upon the above previous *Callithrix* studies, we have conducted the largest to-date geographical sampling of *Callithrix* mitogenomes across Brazil (Figure 1) with the following aims: (1) improve resolution of phylogenetic relationships and divergence times estimates between *Callithrix* mtDNA haplotypes; (2) determine which *Callithrix* mtDNA lineages are autochthonous across *Callithrix* ranges; and (3) identify allochthonous *Callithrix* mtDNA lineages in the southeastern Atlantic Forest and their possible biogeographic origins. We sequenced, for the first time, the complete mitogenome of *C. aurita*, and in total obtained 49 new mitogenome sequences from four species (*C. aurita, C. geoffroyi, C. jacchus, C. penicillata*), and four hybrid types (*C. aurita* x *Callithrix* sp., *C. penicillata* x *C*.*jacchus, Callithrix* sp. x *Callithrix* sp., *C. penicillata* x *C. geoffroyi*) for these analyses.

## Results

Using Illumina whole genome sequencing (WGS) and Sanger sequencing approaches, we sequenced complete mitogenomes from 49 *Callithrix* (Figure 1, Table 1, and Table S1). We combined these new mitogenomes with previously published primate mitogenome sequences for downstream analyses (listed in Table S1). The length of the resulting sequence alignment after combining all of these mitogenomes was 17,132 bases. Sampled individuals that possessed the same mtDNA haplotypes are listed in Table S2. The organization of the *C. aurita* mitogenome was consistent with previously published *Callithrix* mitogenomes from [24]. This mitogenome includes 12 protein-coding genes, two rRNAs, and 14 tRNAs on the heavy strand and one protein-coding gene and eight tRNAs on the light strand, as well as the control region (Table S3). The length of the *C. aurita* mitogenome presented in Table S3 was 16,471 bases.

**Table 1.**
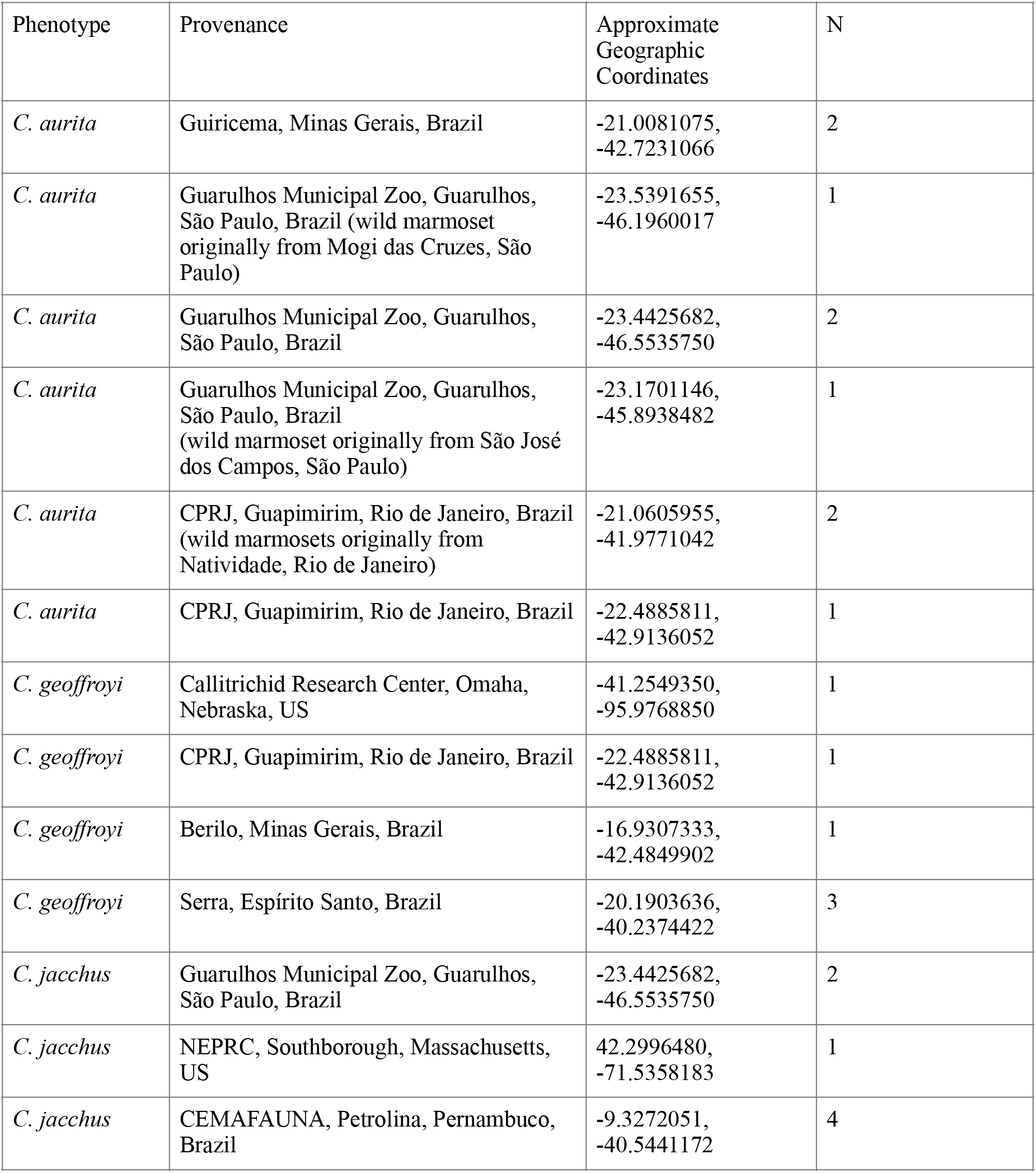
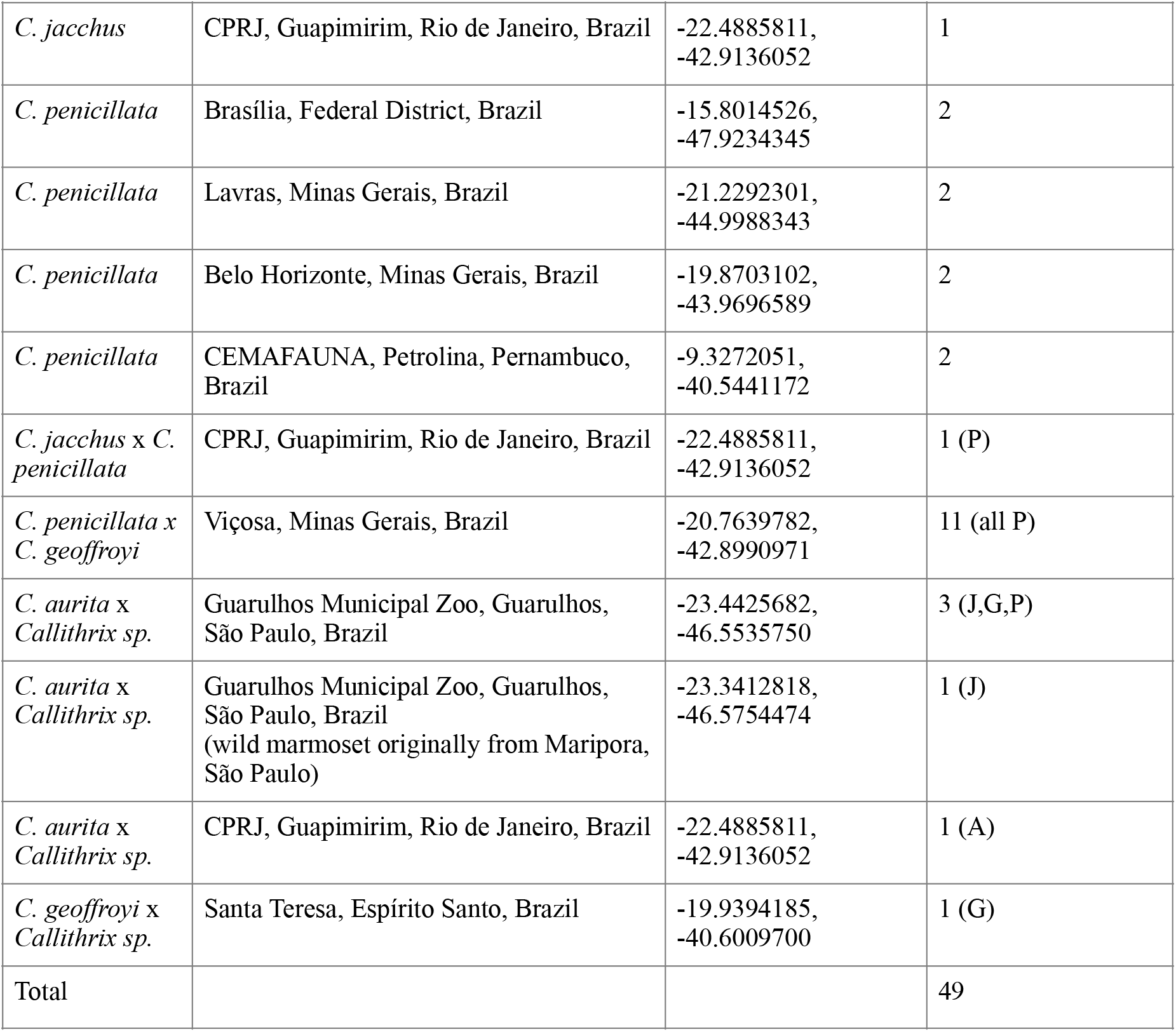
Number of *Callithrix* specimens newly sampled by species and hybrid phenotype. Provenance abbreviations are: CRC-Callitrichid Research Center, NEPRC-New England Primate Research Center (no longer in operation), CPRJ-Centro de Primatologia do Rio de Janeiro, and CEMAFAUNA-Centro de Conservação e Manejo de Fauna da Caatinga. Letters in parentheses next to numerical values listed in the “N” column for hybrid marmosets correspond to likely maternal species of each hybrid based on phylogenetic analyses presented in Figures 2, and S1-S3. Maternal species abbreviations are-A: *C. aurita*, G: *C. geoffroyi*, J: *C. jacchus*; and P: *C. penicillata*.

### Phylogenetic Trees and Divergence Times of Callithrix Mitochondrial Clades

Maximum-likelihood (ML) and Bayesian inference produced well-supported phylogenetic trees that show mostly congruent phylogenetic relationships between the *aurita* and *jacchus* groups (Figure 2, Figures S1-S3). The main difference in the topology of the ML and Bayesian trees was in grouping patterns of some haplotypes within the *C. jacchus* clade described below. A number of nodes in the ML tree possessed 100% bootstrap support but most had bootstrap scores of >70% (Figure S1). Most nodes in the Bayesian trees had posterior probabilities of 1 (Figure 2, Figures S2-S3). Major node names and divergence times within and outside the *Callithrix* clade are shown in Figure 2, Figure S3, Table 2, and Table S4.

**Table 2.** Divergence times in million years (Ma) for *Callithrix* species and select nodes (MRCA = Most recent common ancestor; values in brackets = 95% highest posterior density). Node names follow major node destinations shown in Figure 2 in capital letters.

| Node | Taxa Diverging at Node | Age (Ma) |
| --- | --- | --- |
| A | *C. jacchus* - *C. penicillata* (Caatinga clade) | 0.51 [0.35-0.69] |
| B | *C. kuhlii* - (*C. jacchus* + *C. penicillata* (Caatinga clade)) | 0.82 [0.59-1.09] |
| C | *C. geoffroyi* - (*C. kuhlii* + (*C. jacchus* + *C. penicillata*)) | 1.18 [0.87-1.58] |
| D | *C. aurita* - (*C. geoffroyi* + (*C. kuhlii* + (*C. jacchus* + *C. penicillata*))) | 3.54 [2.37-4.88] |
| E | *Callithrix* - *Cebuella/Mico* | 6.83 [4.86-9.39] |

**Figure 2.**
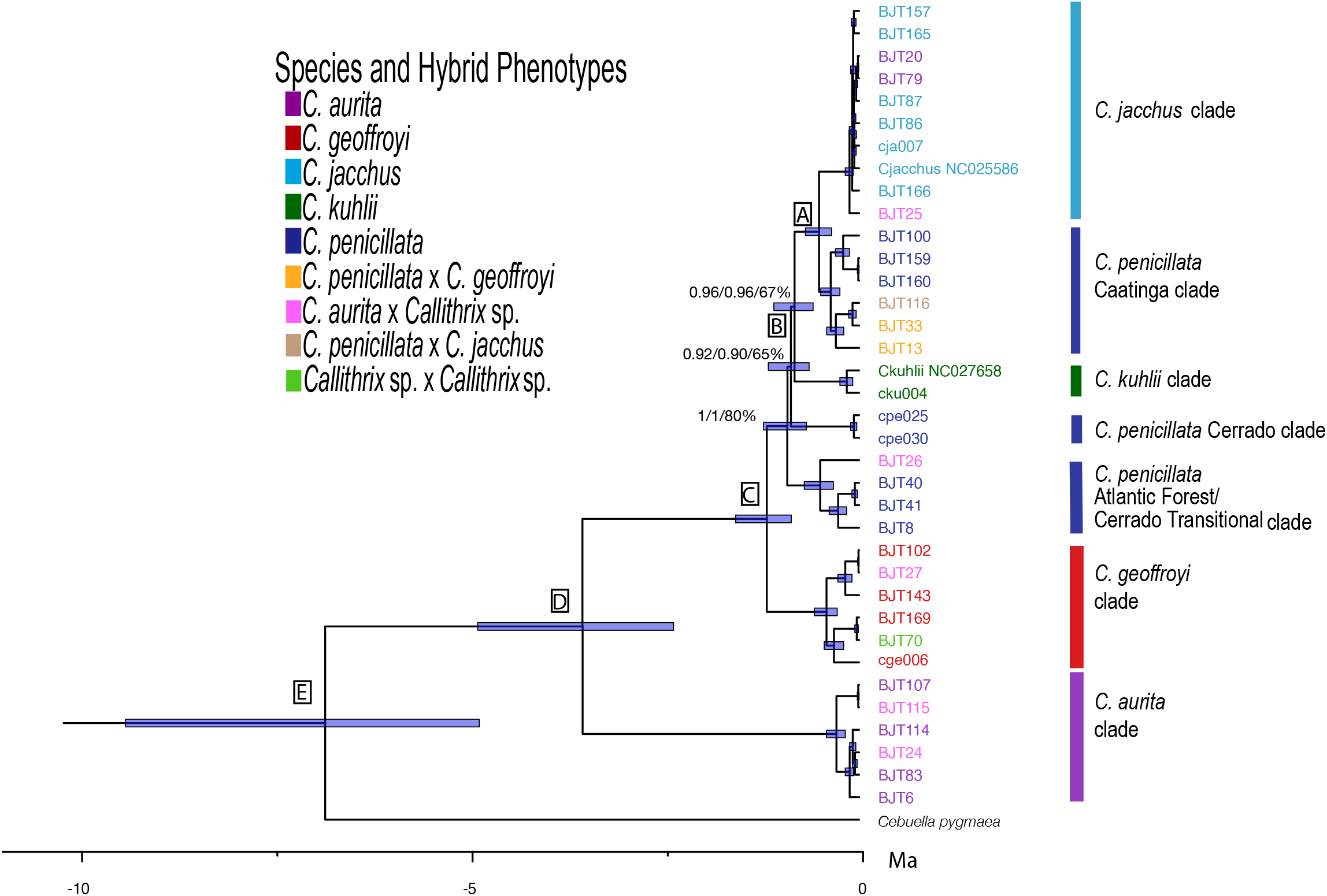
Phylogenetic relationships and divergence ages in million years (Ma) among *Callithrix* haplotypes as calculated from complete mitogenomes (complete tree with outgroups is presented in Figure S3). Major nodes are identified by capital letters, and blue bars at nodes indicate 95% highest posterior densities (HPD) of divergence times. Node support is shown for major nodes where either posterior probability was < 1 in the BEAST tree, posterior probability was < 1 in the MRBAYES tree, or bootstrap support < 70% in the ML tree. Haplotype colors at tips correspond to the ‘Species and Hybrid Phenotypes’ legend, and indicate phenotypes associated with each given haplotype.

*Callithrix* diverged from *Cebuella* approximately 6.83 Ma (Figure 2 node E) and the initial split within *Callithrix*, separating *C. aurita* and the *jacchus* group, occurred approximately 3.54 Ma (Figure 2 node D) (Table 2). Thus, *C. aurita* formed the *Callithrix* basal clade, and *C. geoffroyi* formed the most basal clade within the *jacchus* group by arising 1.18 Ma (node C). *Callithrix penicillata* haplotypes grouped into three polyphyletic clades that corresponded to three different biome regions, an Atlantic Forest-Cerrado transition area, Cerrado, and Caatinga. The first of these *C. penicillata* clades to diverge after *C. geoffroyi* was the Atlantic Forest-Cerrado transition clade at 0.92 Ma. Afterward, the *C. penicillata* Cerrado clade appeared at 0.87 Ma, followed by the *C. kuhlii* clade at 0.82 Ma (Figure 2 node B). The *C. penicillata* Caatinga clade and the *C. jacchus* clades represent the two youngest clades within the phylogeny, splitting about 0.51 Ma (Figure 2 node A). As the *C. jacchus* clade showed some of the shallowest branch tips among *Callithrix* haplotypes and poor phylogenetic resolution, a ParsimonySplits network was constructed for haplotypes within this clade (Figure 3).

**Figure 3.**
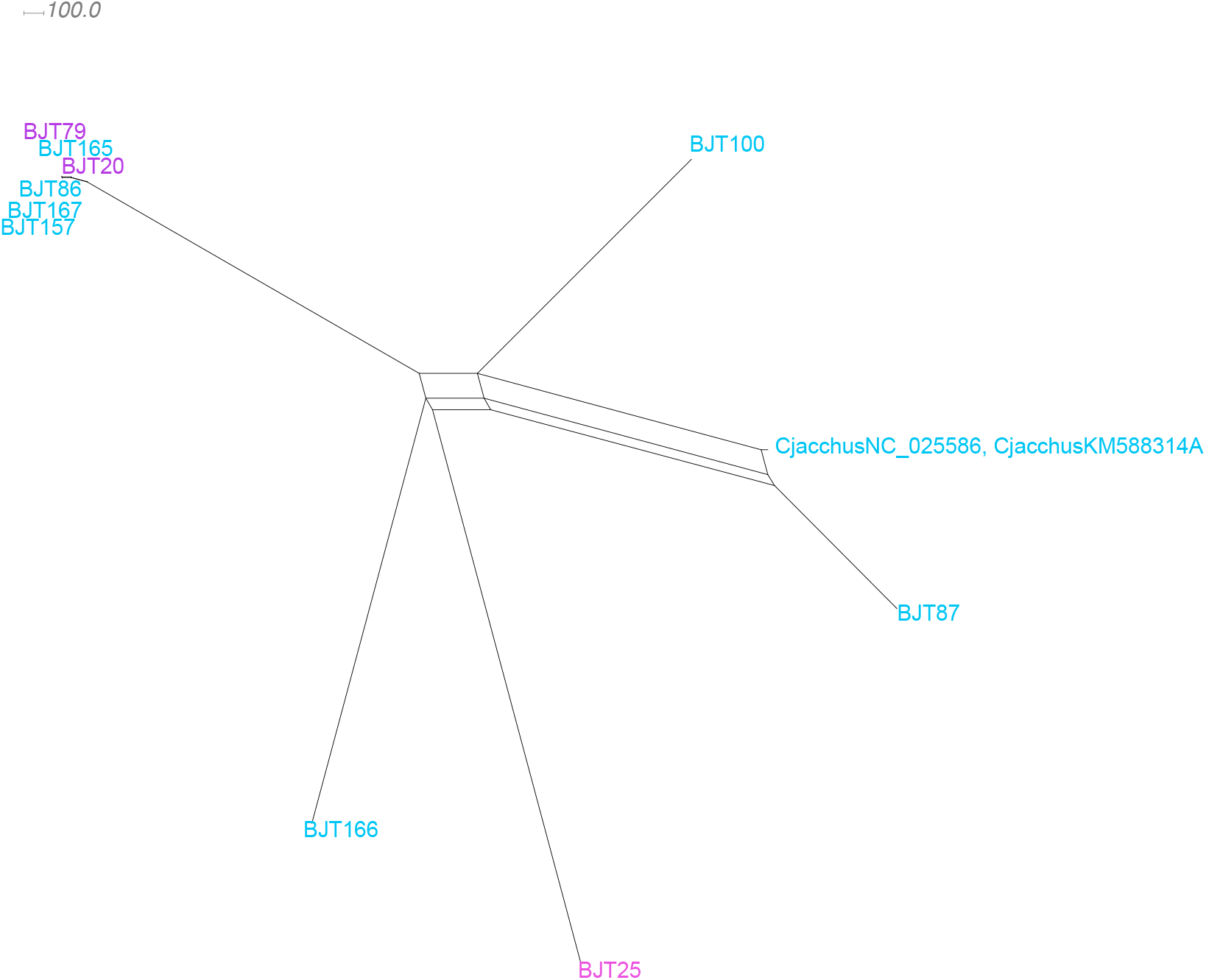
ParimonySplits network of haplotypes from phylogenetic *C. jacchus* clade. Haplotype colors at tips follow Figure 1 ‘Species and Hybrid Phenotypes’ legend, and indicate phenotypes associated with each given haplotype

### Ancestral Origins and Biogeography of Callithrix Mitogenomes

The ancestral origins of *Callithrix* phylogenetic mitogenome clades and subclades based on BMM biogeographic analysis were largely concordant with the assigned Brazilian states and regions of origin of sampled mitogenomic haplotypes (Figure 4 and Table S5). BMM analyses resulted in > 70% posterior probability of an ancestral origin for Node 93, which represented the basal node of the *C. aurita* clade, in Rio de Janeiro state. Within the *C. aurita* clade, node 92 showed >97% posterior probability of an ancestral original of Rio de Janeiro state for two haplotypes sampled within this region from *C. aurita*-phenotype individuals and a *C. aurita* x *Callithrix* sp. hybrid. On the other hand, BMM analysis for nodes 89-91, which represent the other *C. aurita* subclade, assigned posterior probabilities between 44%-65% for an origin of the Minas Gerais state portion of the natural *C. aurita* range. These haplotypes were obtained from *C. aurita*-phenotype individuals sampled in Minas Gerais, São Paulo, and Rio de Janeiro states, as well as a *C. aurita* x *Callithrix* sp. hybrid from São Paulo state.

**Figure 4.**
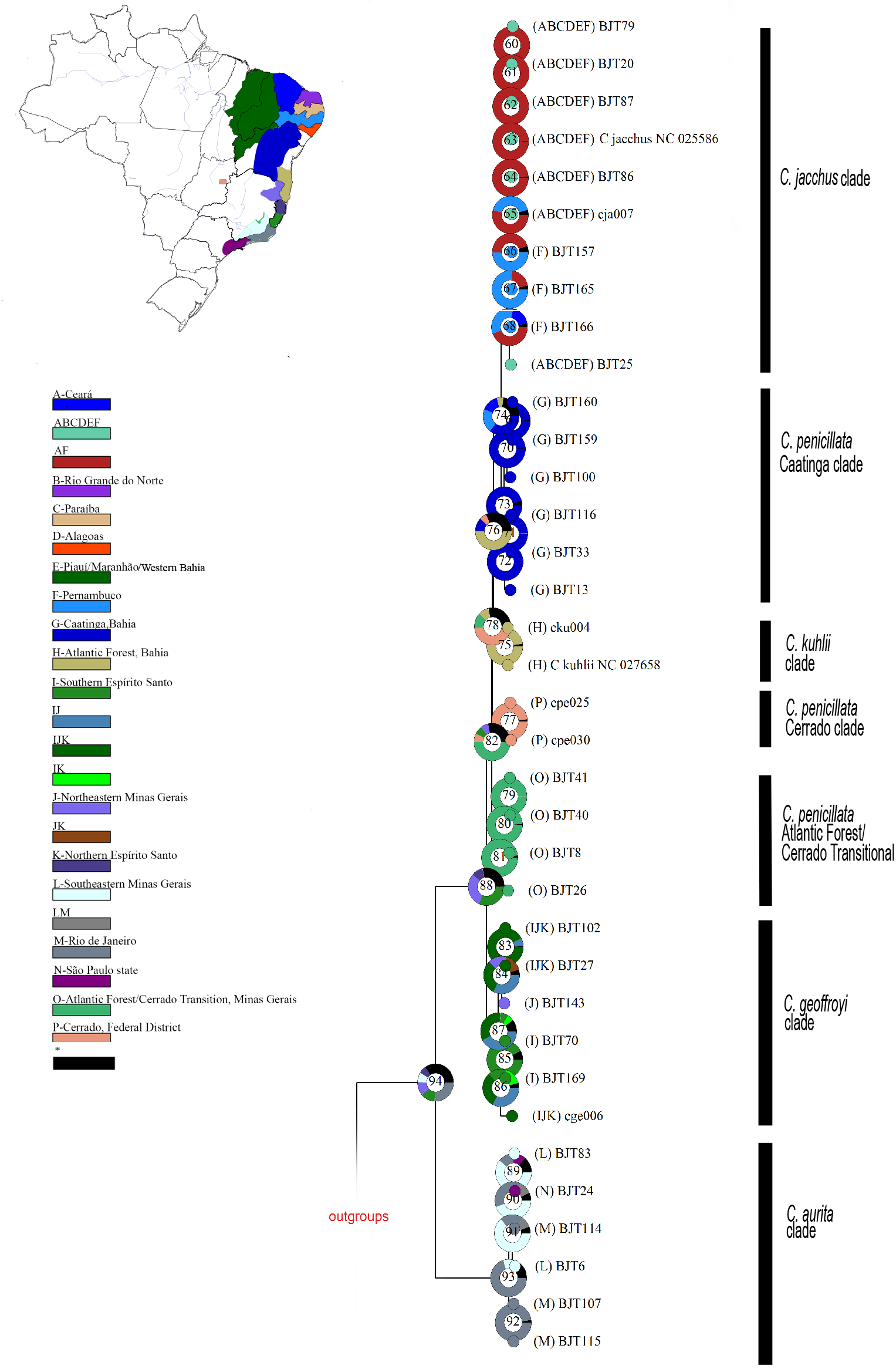
Ancestral state reconstructions performed by the Bayesian Binary MCMC analysis as implemented in RASP v4.2 using the ML rooted tree. Donut charts at each node represent ancestral host estimations. Each node is internally identified with a number. The posterior probabilities of ancestral origins of major nodes are shown in Table S5. Localities where species associated with each phylogenetic clade were sampled or known to occur: A-Ceará state; B-Rio Grande do Norte state; C-Paraíba state; D-Alagoas state; E-Piauí/Maranhão/ Western Bahia states; F-Pernambuco state; G-Caatinga biome in Bahia state; H-Atlantic Forest biome in Bahia state; I-southern Espírito Santo state; J-northeastern Minas Gerais; K-northern Espírito Santo state; L-southeastern Minas Gerais; M-Rio de Janeiro state; N-São Paulo state; O-Atlantic Forest and Cerrado transitional areas in southern Minas Gerais; P-Cerrado Brazilian Federal District. These localities are color coded in the map inset on the left side.

Node 87 in Figure 4 represents the basal node of the *C. geoffroyi* clade, and BMM analyses calculated a collective posterior probability of over 75% of this clade originating within the natural range of *C. geoffroyi*. With a BMM posterior probability of 91.93% that node 85 originated in southeastern Espírito Santo state, biogeographic analysis accurately reflected the sampling origin of haplotypes BJT70 and BJT169. These haplotypes come from *C. geoffroyi-*phenotype individuals, as well as one *Callithrix sp*. x *Callithrix sp*. hybrid. For the other *C. geoffroyi* subclade, BMM analyses posterior probabilities support an ancestral origin of associated haplotypes within the natural distribution of *C. geoffroyi*.

For the three *C. penicillata* clades, BMM analysis showed high posterior probabilities for each clade’s corresponding geographic area as also being each respective clade’s ancestral region. Nodes 79-81 (Figure 4), which represent the *C. penicillata* Atlantic Forest-Cerrado transition clade, each possessed > 98% posterior probabilities of originating in the Atlantic Forest-Cerrado transition zone of Minas Gerais. This clade contained several haplotypes from *C. penicillata*-phenotype individuals sampled in this transition zone, as well as a hybrid sampled in São Paulo state. BMM posterior probability for the central Brazil Cerrado being the ancestral region for node 77 (Figure 4), which encompassed the *C. penicillata* Cerrado clade, was 98.16%. The *C. penicillata* Cerrado clade included haplotypes from *C. penicillata-*phenotype individuals sampled in Brasília. Finally, nodes 69-73 (Figure 4), representing the *C. penicillata* Caatinga clade, possessed BMM posterior probability support between 96.14%-99.60% for the Caatinga of Bahia state as the ancestral region of this clade. The clade contained haplotypes from *C. penicillata-*phenotype animals sampled at CEMAFAUNA. Two haplotypes, representing eleven *C. penicillata* x *C. geoffroyi* hybrids sampled in Viçosa as well as a *C. jacchus* x *C. penicillata* hybrid, clustered within the *C. penicillata* Caatinga group.

BMM biogeographic analysis of the *C. jacchus* clade calculated high posterior probability (>99%) that haplotypes associated with nodes 60-64 originated in Ceará and/or Pernambuco states, regions whose dominant biome is the Caatinga. These haplotypes were obtained from marmosets with *C. jacchus* phenotypes sampled at CEMAFAUNA and the Guarulhos Zoo, as well as three *C. aurita* phenotype individuals sampled within São Paulo. For nodes 65-68, BMM analyses calculated posterior probabilities of the associated haplotypes originating first from Pernambuco, and then from Ceará and/or Pernambuco. In particular, nodes 66 and 67 had respective posterior probabilities of 49.68% and 76.88% of originating in Pernambuco state. Haplotypes associated with these nodes came from a *C. aurita* x *Callithrix* sp. hybrid sampled in São Paulo state and a CEMAFAUNA *C. jacchus-*phenotype individual.

### Genetic Distance between Callithrix Phylogenetic Clades

Pairwise genetic distances between the above established phylogenetic clades are shown in Table 3 as measures of D_xy_. The *C. aurita* clade was the most genetically distant from all other *Callithrix* clades, with D_xy_=0.055-0.056. The smallest genetic distance can be observed between *C. jacchus* and the *C. penicillata* Caatinga clade at D_xy_=0.009. The remaining pairwise genetic distances varied between D_xy_=0.013-0.015, but the *C. geoffroyi* clade was the most distant relative to all other *jacchus* group clades.

**Table 3.** Array of pairwise D_xy_ genetic distances between *Callithrix* phylogenetic clades.

|  |  |  |  |  |  |  |
| --- | --- | --- | --- | --- | --- | --- |
|  | *C. jacchus* |  |  |  |  |  |
| *C. penicillata*<br>(Caatinga clade) | 0.009 | *C. penicillata*<br>(Caatinga clade) |  |  |  |  |
| *C. kuhlii* | 0.014 | 0.013 | *C. kuhlii* |  |  |  |
| *C. penicillata*<br>(Cerrado clade) | 0.016 | 0.015 | 0.014 | *C. penicillata*<br>(Cerrado clade) |  |  |
| *C. penicillata*<br>(Atlantic Forest/<br>Cerrado<br>Transitional<br>clade) | 0.015 | 0.016 | 0.014 | 0.015 | *C. penicillata*<br>(Atlantic<br>Forest/<br>Cerrado<br>Transitional<br>clade) |  |
| *C. geoffroyi* | 0.018 | 0.017 | 0.017 | 0.018 | 0.017 | *C. geoffroyi* |
| *C. aurita* | 0.056 | 0.056 | 0.055 | 0.056 | 0.055 | 0.056 |

## Discussion

### Callithrix *Mitochondrial Phylogenetic Relationships and Divergence Times*

Our ML and Bayesian phylogenies were generally well supported and corroborated *Callithrix* divergence patterns from previous nuclear and mtDNA studies [3,24,25,26]. In ours and these previous phylogenies, the *C. aurita* clade was the most basal within the genus, the *C. geoffroyi* clade was most basal within the *jacchus* group, and *C. penicillata* and *C. jacchus* was the most recently diverged sister clade. Finally, our mtDNA analysis also showed that *C. penicillata* mitochondrial clades are polyphyletic, similar to the results obtained by [21] and [23]. The latter two studies also showed that *C. kuhlii* mitochondrial clades are polyphyletic.

Given the recent divergence times of *Callithrix* species, *Callithrix* polyphyly may be explained by incomplete lineage sorting when ancestral polymorphisms at a given locus are not fixed before population divergence [27]. Another possibility to explain *Callithrix* polyphyly may be due to past hybridization between these species and other *Callithrix* taxa or perhaps recent migrations of *C. kuhlii* outside of its native range. However, we do not believe it is likely for any of these cases. For these alternative scenarios, we would expect to find at least some instances of allochthonous *C. kuhlii* mtDNA lineages, which to our knowledge have not yet been reported. Additionally, we did not observe any discordance involving *C. kuhlii* genotypes/phenotypes with that of any other *Callithrix* species, nor did mitogenome haplotypes from any hybrids sampled in this or previous studies group with *C. kuhlii* phylogenetic clades. Finally, locations where we sampled *Callithrix* species within native ranges were far removed from any natural hybridization zones with *C. kuhlii*, so natural secondary contact between *C. kuhlii* and other *Callithrix* species was unlikely at our sampling locations. Thus, incomplete lineage sorting is the most parsimonious explanation for *Callithrix* polyphyly observed in this and previous *Callithrix* studies.

The *Callithrix* divergence time estimates from our study, being between approximately 0.5 and 6.8 Ma, are within the range of previously published estimates [6,25,26]. These time estimates place the divergence of *Callithrix* species into the Pleistocene. In this epoch, climatic oscillations that may have promoted para- and allopatric speciation in South America, including that of *Callithrix* species, through repeated contractions and expansions of forested refuge [28,29].

### Biogeography Origins of Autochthonous and Allochthonous Callithrix Mitogenomes

Previous phylogenetic and biogeographic analyses support a *Callithrix* origin in the southeastern Brazilian Atlantic Forest and then a northward expansion [3,4,24,25,26]. Our biogeographic analysis of *Callithrix* mitogenome lineages reflected similar biogeographic patterns for *Callithrix* species, as well as a natural geographic separation of major phylogenetic clades. When considering sampled mitogenome lineages of known provenance, the biogeographic origins of our reconstructed phylogenetic clades was strongly influenced by the geographic origin of our samples across natural *Callithrix* ranges. For example, within the *C. geoffroyi* clade, the Minas Gerais state lineage formed a separate clade from the Espírito Santo state lineages. *Callithrix penicillata* possesses the largest natural geographic distribution of all *Callithrix* species [5] and biogeographic origins of paraphyletic clades were defined by where samples were collected within the Cerrado, Atlantic Forest, and Caatinga biomes. Although most sampled *C. jacchus* clade haplotypes did not possess known provenance and showed shallow tips, biogeographic analyses showed strong evidence that these haplotypes likely originated from within the Caatinga biome. Further evidence that *C. jacchus* mtDNA lineages tend to group geographically under denser sampling was shown by [23] with a geographically broader sampling of *C. jacchus* mtDNA D-loop sequences.

Allochthonous *Callithrix* species began appearing in portions of the southeastern Brazilian Atlantic Forest within approximately the last 20-30 years [30,31, pers. obs. C. Igayara]. Our biogeographic analyses can be used to infer the probable origins of parental populations of these allochthonous *Callithrix* species and anthropogenic hybrid *Callithrix* found in the southeastern Brazilian Atlantic Forest. Overall, biogeographic patterns show that the parental populations of these *Callithrix* likely possess multiple geographic origins from within and outside Atlantic Forest. For example, our biogeographic results show that mtDNA haplotypes of three *C. aurita* x *Callithrix* sp. hybrids we sampled in São Paulo state respectively originated from northeastern Minas Gerais or Espírito Santo states, the Atlantic Forest-Cerrado Transitional region, and the Caatinga. On the other hand, the likely provenance of a haplotype of *C. aurita* x *Callithrix* sp. individual was Guapimirim, Rio de Janeiro state. The three haplotypes from *C. penicillata* x *C. geoffroyi* hybrids we sampled in Minas Gerais state and a *C. jacchus* x *C. penicillata* hybrid sampled from Rio de Janeiro state likely originated from the Caatinga.

### Implications of Biological Invasions for Callithrix Genetic Integrity, Hybridization, and Conservation

Several species of non-native fauna and flora have been introduced to the Brazilian Atlantic Forest [32] - one of the most anthropogenically disturbed, yet highly biodiverse biomes on Earth [33,34]. Deliberate and accidental relocations of species beyond their natural geographic ranges by humans may lead to the establishment of non-autochthonous populations and biological invasions within new geographic localities. Such introductions alter the ecological relationships among taxa, and in cases of closely related species, gene flow may occur due to hybridization [35,36,37].

Indeed, our analyses show, for the first time, evidence of introgression of genetic material from allochthonous *Callithrix* species into the genetic background of an endangered, autochthonous *Callithrix* species. Two mtDNA haplotypes that grouped within the *C. jacchus* clade were associated with three individuals with pure *C. aurita* phenotypes sampled within São Paulo state. Two of these individuals were sampled in two different regions of São Paulo state in municipalities (Mogi Das Cruzes and São José dos Campos) that lie 60 km apart. These data not only show the first genetic evidence for cryptic hybridization within the *aurita* group marmosets, but also suggest two independent occurrences of a *C. jacchus* female mating with a *C. aurita* male that led to genetic introgression. Under scenarios of biological invasions, theoretical and empirical data show that hybridization between allochthonous species and endangered, native species creates extinction risk for the latter [38]. Our initial mtDNA data strongly prompt for the development of diagnostic genetic markers to detect the actual extent of allochthonous *Callithrix* genetic introgression in *C. aurita* populations, particularly within São Paulo state.

Contemporary anthropogenic hybrid *Callithrix* and allochthonous *Callithrix* species are normally found in urban or peri-urban areas of southeastern Brazil, due to releases of exotic pet marmosets into such locales [6,9,31,10], where they may encounter autochthonous *Callithrix* species. Indeed, cases exist of native *C. aurita* and *C. flaviceps* meeting up and interbreeding with hybrid and allochthonous *Callithrix* at urban fringes [6,9,10,11,39,40]. Such interactions likely facilitate gene flow from invasive *C. jacchus* and *C. penicillata* into marmoset populations in southeastern Brazil, with consequences that may include outbreeding depression, admixture, hybrid swamping, or introgressive replacement [41,42,43,44].

Invasive *C. jacchus* and *C. penicillata* represent a potential risk for genetic extinction of the other two *jacchus* group species, *C. geoffroyi* and *C. kuhlii*, and [10] recently showed that *C. penicillata* is encroaching on the range of *C. geoffroyi*. We sampled one *Callithrix* sp. x *Callithrix* sp. hybrid in Santa Teresa, Espírito Santo state, a city that straddles the native ranges of *C. flaviceps* and *C. geoffroyi*. Although this hybrid possessed a *C. geoffroyi* clade mitogenome lineage, the individual had a phenotype that strongly suggested some level of *C. penicillata* or *C. jacchus* ancestry-a white “star” on the forehead [30]. Anthropogenic hybridization of *jacchus* group species generally results in the formation of hybrid swarms, admixed populations that lost parental phenotypes and genotypes [6,30,45]. Should large numbers of exotic *C. jacchus* or *C. penicillata* ever invade native ranges of *C. kuhlii* or *C. geoffroyi*, the latter two species may be threatened with genetic swamping by the former two species, a process through which parental lineages are replaced by hybrids that have admixed genetic ancestry [38]. As *C. kuhlii* is considered vulnerable [46], biological invasions by other marmosets present potential conservation risks for this species.

Brazil already possesses several legal instruments for the conservation and protection of wildlife [discussed in 47]. These instruments include national species plans that legally lay out action and aims for the protection of specific groups of endangered species. A national species plan already exists that includes *C. aurita* and *C. flaviceps*, the National Action Plan for the Conservation of Atlantic Forest Primates and Collared Sloth [PAN PPMA, 47], and this plan may eventually need to include *C. kuhlii*. The PAN PPMA considers hybridization as a major threat to the survival of the *aurita* group marmoset species. Thus, the expanded perspective on marmoset hybridization provided by this work should be considered within the context of Brazilian legal instruments that protect endangered marmosets. Such an evaluation is important for incorporating new biological information about marmoset hybridization, as it may call for adopting new legal measures or modifying existing ones to further protect endangered Brazilian primates.

## Conclusions

We provide a robust *Callithrix* phylogeny based on the largest to-date geographical sampling of *Callithrix* mitogenomes across Brazil, showing that the *aurita* group is basal to the *jacchus* group. Our divergence time estimates show these two groups diverged approximately 3.54 Ma, and within the *jacchus* group, *C. jacchus* diverged most recently from the *C. penicillata* Caatinga clade approximately 0.51 Ma. With future sampling of *C. flaviceps*, full mitogenomes can likely be utilized to fully resolve the *Callithrix* phylogeny. Nonetheless, we used our current well-supported phylogenies and biogeographic analyses to elucidate, for the first time, evolutionary relationships among autochthonous, allochthonous, and anthropogenic hybrid marmosets across Brazil. We show that parental populations of allochthonous and anthropogenic hybrid marmosets within the southeastern Brazilian Atlantic Forest incorporate local populations and populations broadly distributed outside of the regions. We also show, for the first time, evidence of allochthonous *Callithrix* species genetic introgression into the genetic background of endangered, autochthonous *C. aurita*. At this time, further determination is needed of the ancestry of *Callithrix* anthropogenic hybrids in southeastern Brazil as well as the fitness and viability of these hybrids. Such data will help determine to what extent anthropogenic hybrids and allochthonous *Callithrix* species threaten the genetic integrity, or ability of a population to preserve its genotypes over generations [48], of autochthonous Atlantic Forest *Callithrix* species.

## Methods

### Samples

In 2011, skin samples were collected from two *C. penicillata* individuals that were captured in Brasília, Federal District. Between 2010 and 2016, skin tissue was collected from: (1) wild marmosets in Minas Gerais and Espírito Santo states as well as the Brazilian Federal District; (2) captive-born, wild-caught, and confiscated marmosets housed at the Guarulhos Municipal Zoo, Guarulhos, São Paulo, CEMAFAUNA (Centro de Manejo de Fauna da Caatinga), Petrolina, Pernambuco, and Centro de Primatologia do Rio de Janeiro (CPRJ), Guapimirim, Rio de Janeiro; (3) a wild group from Natividade, Rio de Janeiro that was caught and housed at CPRJ; (4) a captive-born *C. geoffroyi* sample donated by the Callitrichid Research Center (CRC), Omaha, Nebraska, US; (5) a captive born *C. jacchus* donated by the New England Primate Research Center (NEPRC, now closed), Southborough, Massachusetts, US. Sampling consisted of a total of 49 *Callithrix* individuals as described in Table 1, Table S1, and Figure 1. Table S1 also lists information on utilized sequences that were published elsewhere. Marmoset capture and sampling methodology has been described elsewhere [23]. All individuals were allowed to recover after sample collection, and wild marmosets were released at their point of capture. Specimens were classified phenotypically as pure *C. aurita, C. geoffroyi, C. jacchus* and *C. penicillata* or hybrid (*C. aurita* x *Callithrix* sp., *C. jacchus* x *C. penicillata*, and *C. penicillata* x *C. geoffroyi*) based on published descriptions [2,11,23,30].

### Laboratory Protocols

DNA from skin samples was extracted using a standard proteinase K/phenol/ chloroform protocol [49]. Buffers used for extraction, precipitation and elution of DNA from blood and skin tissue are listed elsewhere [24]. DNA from the Callitrichid Research Center samples was extracted at Arizona State University (ASU). DNA from Brasília individuals was extracted at Northern State Fluminense University, Rio de Janeiro State, Brazil, and then exported to ASU (CITES permit #11BR007015/DF). DNA from all other individuals was extracted at the Federal University of Viçosa (UFV), Viçosa, Minas Gerais, Brazil.

Mitogenomes were obtained for a subset of the samples (Table S1) following the long-range PCR (LR-PCR) methodology of [24], and sequenced on an ABI 3730 sequencer with the BigDye Cycle Sequencing Kit (Applied Biosystems) by the ASU School of Life Science DNA Core Laboratory. The remainder of mitogenomes was obtained from whole genome sequencing (WGS). Individual WGS sequencing libraries were prepared at UFV and ASU with Illumina Nextera DNA Flex Library Prep Kits (catalog #20018704) following manufacturer’s instructions. Individual libraries were barcoded with Illumina Nextera DNA CD Indexes (catalog # 20018707), and pooled in equimolar amounts and sequenced on an Illumina NextSeq using v2 chemistry for 2 x 150 cycles at the ASU Genomic Core Facilities.

### Mitogenome Alignment and Data Analysis

Genetic samples collected since 2015 have been registered in the Brazilian CGen SISGEN database (Supplementary Table S6) and newly sequenced mitogenomes have been deposited in GenBank (Table S1). Trace files of resulting forward and reverse reads from LR-PCR products for each individual sequence were inspected by eye and merged into a single contig for each sampled individual using SEQMAN PRO software from the DNAStar Lasergene Core 10 suite (DNASTAR, Madison, WI). Mitogenomes from WGS data were assembled with NOVOPlasty 2.6.4 [50] (scripts available at https://github.com/Callithrix-omics/Callithrix_mtDNA.git). We downloaded mitogenome sequences of several primate species from GenBank (Supporting Information Table S1). All mitogenomes were aligned in MAAFT (https://www.ebi.ac.uk/Tools/msa/mafft/) with default settings and this MAFFT alignment was confirmed visually in Mesquite 3.5 [51]. Gene, tRNA, rRNA, and control region features within the newly generated marmoset mitogenomes were manually annotated based on the GenBank record of *C. kuhlii* (Accession number KR817257). To check for the presence of nuclear mitochondrial DNA (numts) in mitochondrial sequence data, we followed the strategy described in [24].

We kept mitogenomes in their entirety, but trimmed part of tRNA-Phe, 12s rRNA and the control region to accommodate the length of all utilized sequences. Mitogenome haplotypes were determined with DnaSP 6.12.03 [52], and haplotypes were used for phylogenetic reconstruction. Individuals that possess identical mtDNA haplotypes are listed in Table S3, and these groups are represented in phylogenetic reconstructions by a single haplotype. We added data from several other New World monkeys (Table S1) and reconstructed phylogenetic trees with ML and Bayesian algorithms using IQ-TREE 2.0.3 [53] and MrBayes 3.2.6 [54,55], respectively. For the ML phylogeny, we used the optimal substitution model (GTR+F+R4) as calculated with ModelFinder [56,57] in IQ-TREE under the Bayesian Information Criterion (BIC). We performed the ML analysis in IQ-TREE with 10,000 ultrafast bootstrap (BS) replications [58]. In MrBayes, we used the closest available substitution model GTR+G. The Bayesian tree was reconstructed via Markov Chain Monte Carlo (MCMC) runs with 10,000,000 generations and tree and parameter sampling occurring every 100 generations. Upon completion of the two runs, the first 25% of generations were discarded as burn-in. To check convergence of all parameters and the adequacy of the burn-in, we assessed the uncorrected potential scale reduction factor (PSRF) [59] and that all parameter Estimated Sample Size (ESS) values were above 200. We calculated posterior probabilities (PP) and a phylogram with mean branch lengths from the posterior density of trees using MrBayes. Phylogenetic trees were visualized and edited with FigTree 1.4.2 (http://tree.bio.ed.ac.uk/software/figtree/). Pairwise genetic distances between each of the resulting *Callithrix* mitochondrial clades was measured in DnaSP 6.12.03 as D_xy_, the average number of per site nucleotide substitutions between clades.

The divergence time calculation was performed with the BEAST 2.4.8 package [60] using a relaxed lognormal clock model of lineage variation [61] and by applying a Yule tree prior and the best-fit model of sequence evolution as obtained by ModelFinder. To calibrate the molecular clock, we applied fossil data to constrain the splits between Cebinae and Saimirinae and between Callicebinae and Pitheciinae with hard minimum and soft maximum bounds using a log normal prior following settings and fossils described in detail in [62]. Briefly, for the Cebinae - Saimirinae split, we used an offset of 12.6, mean of 1.287 and standard deviation of 0.8, which translates into a median divergence of 16.2 million years ago (Ma) (95% highest posterior density [HPD]: 13.4-30.0 Ma). For the Callicebinae – Pitheciinae split, we used an offset of 15.7, mean of 1.016 and standard deviation of 0.8, resulting in a median divergence of 18.5 Ma (95% HPD: 16.3-28.9 Ma). We performed two independent runs each with 50 million generations and tree and parameter sampling setting in every 5000 generations. To assess the adequacy of a 10% burn-in and convergence of all parameters, we inspected the trace of the parameters across generations using Tracer 1.6 [63]. We combined sampling distributions of both replicates with LogCombiner 2.4.8 and summarized trees with a 10% burn-in using TreeAnnotator 2.4.8 (both programs are part of the BEAST package). A ParimonySplits network of a subset of mtDNA haplotypes was made with default settings in SplitsTree4 [64].

To reconstruct the biogeographic history of *Callithrix* mitochondrial lineages, we applied the Bayesian Binary Method (BBM) in Reconstruct-Ancestral-States-in-Phylogenies 4.0 (RASP) [65,66]. The ML phylogeny obtained with IQTREE was used for the BBM analysis, which was conducted as two independent runs of 10 chains that ran for 5,000,000 generations and sampled every 100 generations. The fixed Jukes-Cantor+Gamma evolutionary model was implemented for each run. For haplotypes states of origin within a given phylogenetic clade, presence and absence of each associated taxon was determined using a combination of information of known provenance for sampled individuals and recognized *Callithrix* geographical distribution following [5]. For haplotypes obtained from exotic *Callithrix* species and anthropogenic hybrids, we noted where each of these haplotypes clustered among resulting phylogenetic clades, and assigned probable origin for these haplotypes according to the likely natural geographic range associated with each clade.

## Abbreviations

ASU: Arizona State University
BBM: Bayesian Binary Method
BIC: Bayesian Information Criterion
BS: Bootstrap
CEMAFAUNA: Centro de Conservação e Manejo de Fauna da Caatinga
CPRJ: Centro de Primatologia do Rio de Janeiro
CRC: Callitrichid Research Center
ESS: Estimated Sample Size
HPD: Highest posterior density
LR-PCR: long-range PCR
mtDNA: mitochondrial DNA
Ma: million years ago
MCMC: Markov Chain Monte Carlo
MRCA: Most recent common ancestor
ML: Maximum likelihood
NEPRC: New England Primate Research Center
PP: Posterior probabilities
PSRF: potential scale reduction factor
UFV: Universidade Federal de Viçosa
WGS: Whole genome sequencing

## Declarations

### Ethics approval and consent to participate

Tissues were collected under the approval of the ASU Institutional Animal Care and Use Committee Animals (ASU IACUC, protocols #11-1150R, 15-144R), Brazilian Environmental Ministry (SISBIO protocols #47964-2 and #28075-2), and a CPRJ internal review. Biological tissue sampling complied with all institutional, national, and international guidelines.

### Consent for publication

Not applicable.

### Availability of data and materials

Newly sequenced and annotated *Callithrix* mitogenomes have been deposited in NCBI GenBank. Accession numbers for each sequence are given in Table S1.

### Authors’ contributions

JM formulated the idea for the study, collected samples, obtained funding, conducted wet and dry laboratory work, and wrote the original manuscript. RAC provided study guidance, and logistical support. NHAC provided field assistance and logistical support. JAD provided study guidance, and logistical support. CM assisted in phenotypic identification of hybrids and generation of Appendix A. CSI gave access and provided logistical support to collect samples from animals kept at Guarulhos Zoo. SBM provided logistical support and veterinary assistance to collect samples from animals kept at the Rio de Janeiro Primatology Center. AN gave assistance with data processing. PAN gave access and provided logistical support to collect samples from animals kept at CEMAFAUNA. MP provided field assistance and logistical support. LCMP gave access and provided logistical support to collect samples from animals kept at CEMAFAUNA. AP gave access and provided logistical support to collect samples from animals kept at CPRJ. CRRM and DZ were major contributors in writing the manuscript. DLS performed DNA extractions, methodological optimization, and carried out PCR. ACS was a major contributor in writing the manuscript, provided study guidance, and logistical support. CR gave assistance in data processing, carried out divergence analysis, gave significant guidance and input in the development of this study. All authors read and approved the final manuscript.

## Acknowledgements

We would like to thank the Callitrichid Research Center, the Golden Lion Tamarin Association, CEMAFAUNA, Rio de Janeiro Primatology Center, the Guarulhos Municipal Zoo, the Beagle Lab at UFV, and many biologists, field technicians, veterinarians, and other individuals that made this research possible. We would like to thank two anonymous reviewers for insightful comments on a previous version of this work.

## References

1. Finstermeier K, Zinner D, Brameier M, Meyer M, Kreuz E, Hofreiter M, Roos C. A mitogenomic phylogeny of living primates. PLoS ONE. 2013;8:e69504.

2. Malukiewicz J, Boere V, Fuzessy LF, Grativol AD, De Oliveira e Silva I, Pereira LCM, et al. Natural and anthropogenic hybridization in two species of eastern Brazilian marmosets (Callithrix jacchus and C. penicillata). PLoS ONE. 2015; 10(6):e0127268.

3. Perelman P, Johnson WE, Roos C, Seuanez HN, Hovarth JE, Moreira AM, et al. A molecular phylogeny of living primates. PLoS Genet. 2011; 7:e1001342.

4. Buckner JC, Lynch-Alfaro JW, Rylands AB, Alfaro ME. Biogeography of the marmosets and tamarins (Callitrichidae). Mol Phylogenet Evol. 2015; 82B:413–425.

5. Rylands AB, Coimbra-Filho AF, Mittermeier RA. The systematics and distributions of the marmosets (Callithrix, Callibella, Cebuella, and Mico) and Callimico (Callimico) (Callitrichidae, Primates). In: Ford SM, Porter LM, Davis LC, editors. The smallest anthropoids: the marmoset/Callimico radiation. New York: Springer; 2009. p. 25–61

6. Malukiewicz J. A Review of experimental, natural, and anthropogenic hybridization in Callithrix marmosets. Int J Primatol. 2019; 40:72–98.

7. Moraes AM, Vancine MH, Moraes AM, de Oliveira Cordeiro CL, Pinto MP, Lima AA, et al. Predicting the potential hybridization zones between native and invasive marmosets within Neotropical biodiversity hotspots. Global Ecol Conserv. 2019; 20:e00706.

8. Vale CA, Neto LM, Prezoto F. Distribution and invasive potential of the black-tufted marmoset Callithrix penicillata in the Brazilian territory. Scientia Plena. 2020:052401.

9. Carvalho RS, Bergallo HG, Cronemberger C, Guimarães-Luiz T, Igayara-Souza CA, Jerusalinsky L, et al. Callithrix aurita: A tiny primate on the edge of extinction in the Brazilian Atlantic Forest. Neotropical Primates. 2018; 24:1–8.

10. Silva FFR, Malukiewicz J, Silva LC, Carvalho RS, Ruiz-Miranda CR, Coelho FAS et al. A survey of wild and introduced marmosets (Callithrix: Callitrichidae) in the southern and eastern portions of the state of Minas Gerais, Brazil. Primate Conserv. 2018; 32:1–18.

11. Carvalho RS. Conservação do saguis-da-serra-escuro (Callithrix aurita (Primates)) – Analise molecular e colormetrica de populações do gênero Callithrix e seus híbridos. PhD dissertation, Universidade do Estado do Rio de Janeiro. 2015.

12. Melo F, Bicca-Marques J, Ferraz DS, Jerusalinsky L, Mittermeier RA, Oliveira L, et al. Callithrix aurita. The IUCN Red List of Threatened Species 2019: e.T3570A17936433. https://dx.doi.org/10.2305/IUCN.UK.2019-3.RLTS.T3570A17936433.en. Downloaded on 13 February 2020.

13. Rylands AB, Ferrari SF, Mendes SL. Callithrix flaviceps. The IUCN Red List of Threatened Species 2008: e.T3571A9951402. https://dx.doi.org/10.2305/IUCN.UK.2008.RLTS.T3571A9951402.en. Downloaded on 13 February 2020.

14. Zinner D, Arnold ML, Roos C. The strange blood: Natural hybridization in primates. Evol Anthropol. 2011;20(3):96–103.

15. Zinner D, Wertheimer J, Liedigk R, Groeneveld LF, Christian Roos C. Baboon phylogeny as inferred from complete mitochondrial genomes. Am J Phys Anthropol. 2013; 150:133–140.

16. Chown SL, Hodgins KA, Griffin PC, Oakeshott JG, Byrne M, Hoffmann AA. Biological invasions, climate change and genomics. Evol Appl; 2015:8: 23–46.

17. Castro JA, Picornell A, Ramon M. Mitochondrial DNA: a tool for population genetics studies. Int Microbiol. 1998;1:327–332.

18. Brown WM, George M Jr, Wilson AC. Rapid evolution of animal mitochondrial DNA. Proc Natl Acad Sci USA. 1979; 76:1967–1971.

19. Mundy NI, Pissinatti A, Woodruff DS. Multiple nuclear insertions of mitochondrial cytochrome b sequences in callitrichine primates. Mol Biol Evol. 2000;17:1075– 1080.

20. Schneider H, Bernardi JAR, da Cunha DB, Tagliaro CH,Vallinoto M, Ferrari SF, Sampaio I. A molecular analysis of the evolutionary relationships in the Callitrichinae with emphasis on the position of the dwarf marmoset. ZoolScr: 2012;41:1–10.

21. Tagliaro CH, Schneider MPC, Schneider H, Sampaio IC, Stanhope MJ. Marmoset phylogenetics, conservation perspectives, and evolution of the mtDNA control region. Mol Biol Evol. 1997;14:674–684.

22. Tagliaro CH, Schneider MPC, Schneider H, Sampaio I, Stanhope M. Molecular studies of Callithrix pygmaea (primates, Platyrrhini) based on transferrin intronic and ND1 regions: implications for taxonomy and conservation. Genet Mol Biol. 2000; 23:729–737.

23. Malukiewicz J, Boere V, Fuzessy LF, Grativol AD, French JA, De Oliveira e Silva I, et al. Hybridization effects and genetic diversity of the common and black-tufted marmoset (Callithrix jacchus and Callithrix penicillata) mitochondrial control region. Am J Phys Anthropol. 2014;155(4):522–536.

24. Malukiewicz J, Hepp CM, Guschanski K, Stone AC. Phylogeny of the jacchus group of Callithrix marmosets based on complete mitochondrial genomes. Am J Phys Anthropol. 2017; 162(1):157–169.

25. Dos Reis M, Gunnell GF, Barba-Montoya J, Wilkins A, Yang Z, Yoder AD. Using phylogenomic data to explore the effects of relaxed clocks and calibration strategies on divergence time estimation: primates as a test case. Syst Biol. 2018; 67:594–615.

26. Springer MS, Meredith RW, Gatesy J, Emerling CA, Park J, Rabosky DL et al. Macroevolutionary dynamics and historical biogeography of primate diversification inferred from a species supermatrix. PLoS ONE. 2012;7:e49521.

27. Rogers J, Gibbs RA. Applications of next generation sequencing primate genomics: Emerging patterns of genome content and dynamics. Nat Rev Genet. 2014; 15:347– 359.

28. Kinzey WG. Distribution of primates and forest refuges. In: Prance GT, editor. Biological diversification in the tropics. New York, NY: Columbia University Press. 1982. p 455–482.

29. Turchetto-Zolet AC, Pinheiro F, Salgueiro F, Palma-Silva C. Phylogeographical patterns shed light on evolutionary process in South America. Mol Ecol. 2013; 22:1193–1213.

30. Fuzessy LF, Silva IO, Malukiewicz J, Silva FFR, Pônzio MC, Boere V, Ackermann RR. Morphological variation in wild marmosets (Callithrix penicillata and C. geoffroyi) and their hybrids. Evol Biol. 2014; 41(3):480–493.

31. Ruiz-Miranda CR, Affonso AG, Martins A, Beck B. Distribuição do sagui (Callithrix jacchus) nas areas de ocorrência do mico-leão-dourado (Leontopithecus rosalia) no estado do Rio de Janeiro. Neotropical Primates. 2000;8:98–100.

32. Frehse FA, Braga RR, Nocera GA, Vitale JRS. Non-native species and invasion biology in a megadiverse country: scientometric analysis and ecological interactions in Brazil. Biol Invasions. 2016:18, 3713–3725.

33. Ribeiro MC, Metzger JP, Martensen AC, Ponzoni FJ, Hirota MM. The Brazilian Atlantic Forest: How much is left, and how is the remaining forest distributed? Implications for conservation. Biol Conserv. 2009; 142:1141–1153.

34. Ribeiro MC, Martensen AC, Metzger JP, Tabarelli M, Scarano F, Fortin MJ. The Brazilian Atlantic Forest: a shrinking biodiversity hotspot. In: Zachos F, Habel J, editors. Biodiversity Hotspots. Berlin, Heidelberg: Springer; 2011. p. 5–21.

35. Cortés-Ortiz L, Roos C, Zinner D. Introduction to special issue on primate hybridization and hybrid zones. Int J Primatol. 2019; 40(1):1–8.

36. Crispo E, Moore J, Lee-Yaw J, Gray S, Haller B. Broken barriers: Human-induced changes to gene flow and introgression in animals. BioEssays. 2011; 33(7):508–518.

37. Oliveira LC, Grelle CEV. Introduced primate species of an Atlantic Forest region in Brazil: present and future implications for the native fauna. TCS. 2012;5(1):112–120.

38. Todesco M, Pascual M, Owens G, Ostevik K, Moyers B, Hübner S et al. Hybridization and extinction. Evol Appl. 2016; 9(7):892–908.

39. Detogne N, Ferreguetti AC, Mello JFF, Santana MC, Dias AC, NCJ da Mota, et al. Spatial distribution of buffy-tufted-ear (Callithrix aurita) and invasive marmosets (Callithrix spp.) in a tropical rainforest reserve in southwestern Brazil. Am J Primatol. 2017; 79:e22718.

40. Detogne N. O sagui-da-serra-escuro (Callithrix aurita) e os saguis invasores no Parque Nacional da Serra dos Órgãos, RJ, Brasil: distribuição espacial e estratégias de conservação. Master ‘s thesis, Universidade do Estado do Rio de Janeiro. 2015.

41. Allendorf F, Leary R, Spruell P, Wenburg, J. The problems with hybrids: Setting conservation guidelines. TREE. 2001;16:613–622.

42. McFarlane SE, Pemberton JM. Detecting the true extent of introgression during anthropogenic hybridization. Trends Ecol Evol. 2019;34(4):315–326.

43. Rhymer JM, Simberloff D. Extinction by hybridization and introgression. Annu Rev Ecol Syst 1996; 27:83–109.

44. Wolf DE, Takebayashi N, Rieseberg LH. Predicting the risk of extinction through hybridization. Conserv Biol. 2001;15(4):1039–1053.

45. Malukiewicz J, Boere V, Fuzessy LF, Grativol AD, De Oliveira e Silva I, Pereira LCM, et al. Natural and anthropogenic hybridization in two species of eastern Brazilian marmosets (Callithrix jacchus and C. penicillata). PLoS ONE. 2015; 10(6):e0127268.

46. Neves L, Bicca-Marques J, Jerusalinsky L, Mittermeier RA, Pereira DG, Rylands AB. 2019. Callithrix kuhlii. The IUCN Red List of Threatened Species 2019:e.T3575A17936243. https://dx.doi.org/10.2305/ IUCN.UK.2019-3.RLTS.T3575A17936243.en. Downloaded on 30 March 2020.

47. Malukiewicz J, Boere V, Borstelmann de Oliveira MA, D’Arc M, Ferreira JVA, French J, Houman G, CAI de Souza, Jerusalinsky L, de Melo FR; SB Valença-Montenegro MM.; Moreira, Silva IO, Pacheco FS.; Rogers J, Pissinatti A, del Rosario R, Ross C, Ruiz-Miranda CR, Pereira LCM, Schiel N, da Silva, FFR, Souto A, Šlipogor V, Tardif S. An Introduction to the Callithrix Genus and Overview of Recent Advances in Marmoset Research. Preprints 2020; 2020110256.

48. Tauer C, Stewart JF, Rodney W, Lilly CJ, Guldin JM, Nelson CD. Hybridization leads to loss of genetic integrity in shortleaf pine: unexpected consequences of pine management and fire suppression. J For. 2012:110, 216–224.

49. Sambrook J, Russel DW. Molecular cloning, 3rd ed. Cold Spring Harbor: CSHL Press. 2001.

50. Dierckxsens N, Mardulyn P, Smits G. NOVOPlasty: de novo assembly of organelle genomes from whole genome data. Nucleic Acids Res. 2017; 45:e18.

51. Maddison WP, Maddison D. Mesquite: a modular system for evolutionary analysis. Version 3.51. 2018. http://www.mesquiteproject.org. Accessed 17 Dec 2019.

52. Rozas J, Ferrer-Mata A, JC Sánchez-DelBarrio, Guirao-Rico S, Librado P, Ramos-Onsins SE et al. DnaSP 6: DNA sequence polymorphism analysis of large datasets. Mol Biol Evol. 2017; 34:3299–3302.

53. Nguyen LT, Schmidt HA, Haeseler A, Minh BQ. IQ-TREE: a fast and effective stochastic algorithm for estimating maximum-likelihood phylogenies. Mol Biol Evol. 2015; 32:268–274.

54. Huelsenbeck JP, F Ronquist.. MRBAYES: Bayesian inference of phylogeny. Bioinformatics. 2001;17:754–755.

55. Ronquist F, Huelsenbeck JP. MRBAYES 3: Bayesian phylogenetic inference under mixed models. Bioinformatics. 2003; 19:1572–1574.

56. Chernomor O, von Haeseler A, Minh BQ. Terrace aware data structure for phylogenomic inference from supermatrices. Syst Biol. 2016; 65:997–1008.

57. Kalyaanamoorthy S, Minh BQ, Wong TKF, von Haeseler A, Jermiin LS. ModelFinder: fast model selection for accurate phylogenetic estimates. Nat Methods. 2017; 14:587–589.

58. Minh BQ, Nguyen MA, von Haeseler A. Ultrafast approximation for phylogenetic bootstrap. Mol Biol Evol. 2013; 30:1188–1195.

59. Gelman A, Rubin DB. Inference from iterative simulation using multiple sequences (with discussion). Stat Sci. 1992; 7:457–472.

60. Bouckaert R, Heled J, Kühnert D, Vaughan T, Wu CH, Xie D, et al. BEAST 2: A Software Platform for Bayesian Evolutionary Analysis. PLoS Comp Biol. 2014; 10:e1003537.

61. Drummond AJ, Ho SYW, Phillips MJ, Rambaut A. Relaxed phylogenetics and dating with confidence. PLoS Biol. 2006; 4(5):e88.

62. Byrne H, Rylands AB, Carneiro JC, Lynch JWA, Bertulo F, Silva MNF et al. Phylogenetic relationships of the New World titi monkeys (Callicebus): first appraisal of taxonomy based on molecular evidence. Front Zool. 2016; 13:10.

63. Rambaut A, Suchard MA, Xie D, Drummond AJ. Tracer v1.6. 2014. http://beast.bio.ed.ac.uk/Tracer. Accessed 17 Dec 2019.

64. Huson HD, Bryant D. Application of phylogenetic networks in evolutionary studies. Mol. Biol. Evol. 2006; 23(2):254–267.

65. Yu Y, Blair C, He XJ. RASP 4: Ancestral State Reconstruction Tool for Multiple Genes and Characters. Mol Biol Evol. 2020;37(2):604–606.

66. Yu Y, Harris AJ, Blair C, He XJ. RASP (Reconstruct Ancestral State in Phylogenies): a tool for historical biogeography. Mol Phylogenet Evol. 2015;87:46–49.

