## Supplemental Tables and Figures for "Mitogenomic Phylogeny of *Callithrix* with Special Focus on Human Transferred Taxa"

**Supplement Material**

Table S1. Metadata for newly collected samples as well as primate mitochondrial genome sequences obtained from Genbank. The 'Sample' column gives ID of each new sampled individual or the species for sequences obtained from previous studies. The 'Accession' column gives Genbank accession numbers for each sequence, the 'Assembly Method This Study' column states the manner in which new *Callithrix* mitochondrial genomes were sequenced and assembled (S = Sanger; N = NOVOPlasty2.6.4). The 'Phenotype' column indicates whether the sampled individual possessed a pure species or hybrid phenotype, and capital letters in parentheses next to *Callithrix penicillata* x *Callithrix geoffroyi* category are specific phenotype classifications following Figure 5 in Fuzessy et al. (2014). The 'mtDNA Genome Lineage' column indicates phylogenetic classification of the mitochondrial genome of the sampled individual. The 'Sampling Location' column indicates where each individual was sampled. Nearest cities are located for individuals sampled from the wild, and facilities are indicated for individuals sampled in captivity. The Guarulhos Municipal Zoo is located in Guarulhos, São Paulo, Brazil; CRC (Callitrichid Research Center) is located in Omaha, Nebraska, US; NEPRC (New England Primate Research Center, no longer in operation) was located in Southborough, Massachusetts, US; CPRJ (Centro de Primatologia do Rio de Janeiro) is located in Guapimirim, Rio de Janeiro, Brazil; CEMAFAUNA (Centro de Concervação e Manejo de Fauna da Caatinga) is located in Petrolina, Pernambuco. Abbreviations for Brazilian states in the 'Sampling Location' column are as follows: Espírito Santo (ES), Minas Gerais (MG), Rio de Janeiro (RJ), São Paulo (SP). DF is the Brazilian Federal District. (NA = No data Available).

| Sample | Accession Number | Assembly method | Phenotype | mtDNA lineage | Sampling location | Latitude/ Longitude |
|---|---|---|---|---|---|---|
| BJT6 | MN787074 | N | *C. aurita* | *C. aurita* | Guiricema, MG | -21.010, -42.720 |
| BJT20 | MN787075 | S | *C. aurita* | *C. jacchus* | Mogi das Cruzes, SP (Removed from wild and housed at Guarulhos Municipal Zoo) | -23.523, -46.179 |
| BJT65 | MT041703 | N | *C. aurita* | *C. aurita* | Guiricema, MG | -21.010, -42.720 |
| BJT79 | MN787077 | S | *C. aurita* | *C. jacchus* | Guarulhos Municipal Zoo | NA |
| BJT83 | MN787078 | N | *C. aurita* | *C. aurita* | Sao Jose dos Campos, SP (Removed from wild and housed at Guarulhos Municipal Zoo) | -23.070, -45.933 |
| BJT84 | MN787079 | S | *C. aurita* | *C. jacchus* | Guarulhos Municipal Zoo | NA |

| | | | | | | |
|---|---|---|---|---|---|---|
| BJT107 | MN787080 | N | *C. aurita* | *C. aurita* | Natividade, RJ (Removed from wild and housed at CPRJ) | -21.043, -41.978 |
| BJT109 | MN787081 | N | *C. aurita* | *C. aurita* | Natividade, RJ (Removed from wild and housed at CPRJ) | -21.043, -41.978 |
| BJT114 | MN787082 | N | *C. aurita* | *C. aurita* | Guapirim, RJ (Removed from wild and housed at CPRJ) | -22.488, -42.912 |
| Cgeoffroyi_NC_021 | NC_021941 | NA | *C. geoffroyi* | *C. geoffroyi* | NA | NA |
| cgeo006 | LR745201 | S | *C. geoffroyi* | *C. geoffroyi* | CRC | NA |
| BJT102 | MN787083 | N | *C. geoffroyi* | *C. geoffroyi* | CPRJ | NA |
| BJT143 | MN787084 | N | *C. geoffroyi* | *C. geoffroyi* | Berilo, MG | -16.953, -42.466 |
| BJT169 | MN787086 | N | *C. geoffroyi* | *C. geoffroyi* | Serra, ES | -20.213, -40.267 |
| BJT170 | MN787085 | N | *C. geoffroyi* | *C. geoffroyi* | Serra, ES | -20.213, -40.267 |
| BJT171 | MN787087 | N | *C. geoffroyi* | *C. geoffroyi* | Serra, ES | -20.213, -40.267 |
| cja007 | LR745200 | S | *C. jacchus* | *C. jacchus* | NEPRC | NA |
| Cjacchus KM588314A | KM588314 | NA | *C. jacchus* | *C. jacchus* | NA | NA |
| Cjacchus NC_025586 | NC_025586 | NA | *C. jacchus* | *C. jacchus* | NA | NA |
| BJT86 | MN787088 | N | *C. jacchus* | *C. jacchus* | Guarulhos Municipal Zoo | NA |
| BJT87 | MN787089 | N | *C. jacchus* | *C. jacchus* | Guarulhos Municipal Zoo | NA |
| BJT100 | MN787090 | N | *C. jacchus* | *C. jacchus* | CPRJ | NA |
| BJT157 | MN787091 | N | *C. jacchus* | *C. jacchus* | CEMAFAUNA | NA |
| BJT165 | MN787092 | N | *C. jacchus* | *C. jacchus* | CEMAFAUNA | NA |
| BJT166 | MN787093 | N | *C. jacchus* | *C. jacchus* | CEMAFAUNA | NA |
| BJT167 | MN787094 | N | *C. jacchus* | *C. jacchus* | CEMAFAUNA | NA |
| cpe025 | MN787101 | N | *C. penicillata* | *C. penicillata* | SWPW Quadra 15, Conjunto 5, Brasília, DF | -15.911, -47.953 |
| cpe030 | MN787102 | N | *C. penicillata* | *C. penicillata* | Jardim Botânico, Brasília, DF | -15.861, -47.829 |
| BJT8 | MN787095 | N | *C. penicillata* | *C. penicillata* | Lavras, MG | -21.227, -44.980 |
| BJT10 | MN787096 | N | *C. penicillata* | *C. penicillata* | Lavras, MG | -21.227, -44.980 |
| BJT40 | MN787097 | N | *C. penicillata* | *C. penicillata* | Belo Horizonte, MG | -19.921, -43.990 |
| BJT41 | MN787098 | N | *C. penicillata* | *C. penicillata* | Belo Horizonte, MG | -19.921, -43.990 |
| BJT159 | MN787099 | N | *C. penicillata* | *C. penicillata* | CEMAFAUNA | NA |
| BJT160 | MN787100 | N | *C. penicillata* | *C. penicillata* | CEMAFAUNA | NA |
| cku004 | KR817257 | N | *C. kuhlii* | *C. kuhlii* | CRC | NA |
| Ckuhlili KR869628 | KR869628 | NA | *C. kuhlii* | *C. kuhlii* | NA | NA |
| Ckuhlii NC_027658 | NC_027658 | NA | *C. kuhlii* | *C. kuhlii* | NA | NA |

| | | | | | | |
|---|---|---|---|---|---|---|
| BJT24 | MN787103 | N | *C. aurita* x sp. | *C. aurita* | Guarulhos Municipal Zoo (apprehended animal) | NA |
| BJT25 | MN787104 | N | *C. aurita* x sp. | *C. jacchus* | Guarulhos Municipal Zoo (apprehended animal) | NA |
| BJT26 | MN787105 | N | *C. aurita* x sp. | *C. penicillata* | Guarulhos Municipal Zoo (apprehended animal) | NA |
| BJT27 | MN787106 | N | *C. aurita* x sp. | *C. geoffroyi* | Guarulhos Municipal Zoo (apprehended animal) | NA |
| BJT115 | MN787107 | N | *C. aurita* x sp. | *C. aurita* | Guapirim, RJ | -22.488, -42.912 |
| BJT116 | MN787108 | N | *C. penicillata* x *C. jacchus* | *C. penicillata* | Guapirim, RJ | -22.488, -42.912 |
| BJT70 | MN787109 | N | *Callithrix* sp. x *Callithrix* sp. | *C. geoffroyi* | Santa Teresa, ES | -19.935, -40.596 |
| BJT13 | MN787110 | N | *C. penicillata* x *C. geoffroyi* (D) | *C. penicillata* | Viçosa, MG | -20.755, -42.872 |
| BJT14 | MN787120 | N | *C. penicillata* x *C. geoffroyi* (B) | *C. penicillata* | Viçosa, MG | -20.755, -42.872 |
| BJT15 | MN787117 | N | *C. penicillata* x *C. geoffroyi* (D) | *C. penicillata* | Viçosa, MG | -20.755, -42.872 |
| BJT16 | MN787114 | N | *C. penicillata* x *C. geoffroyi* | *C. penicillata* | Viçosa, MG | -20.755, -42.872 |
| BJT31 | MN787111 | N | *C. penicillata* x *C. geoffroyi* (C) | *C. penicillata* | Viçosa, MG | -20.758, -42.860 |
| BJT33 | MN787112 | N | *C. penicillata* x *C. geoffroyi* (B) | *C. penicillata* | Viçosa, MG | -20.778, -42.863 |
| BJT38 | MN787113 | N | *C. penicillata* x *C. geoffroyi* (A) | *C. penicillata* | Viçosa, MG | -20.755, -42.857 |
| BJT39 | MN787116 | N | *C. penicillata* x *C. geoffroyi* (D) | *C. penicillata* | Viçosa, MG | -20.755, -42.857 |
| BJT75 | MN787115 | N | *C. penicillata* x *C. geoffroyi* | *C. penicillata* | Viçosa, MG | -20.759, -42.866 |
| BJT150 | MN787118 | N | *C. penicillata* x *C. geoffroyi* (A) | *C. penicillata* | Viçosa, MG | -20.759, -42.866 |
| BJT151 | MN787119 | N | *C. penicillata* x *C. geoffroyi* (A) | *C. penicillata* | Viçosa, MG | -20.759, -42.866 |
| *Ateles belzebuth* | KC757386 | NA | NA | NA | NA | NA |
| *Alouatta caraya* | KC757384 | NA | NA | NA | NA | NA |
| *Aotus azarai* | KC757385 | NA | NA | NA | NA | NA |
| *Aotus lemurinus* | FJ785421 | NA | NA | NA | NA | NA |
| *Aotus nancymaae* | NC_018116 | NA | NA | NA | NA | NA |
| *Brachyteles arachnoides* | JX262672 | NA | NA | NA | NA | NA |
| *Cacajao calvus* | KC959985 | NA | NA | NA | NA | NA |

| | | | | | | |
|---|---|---|---|---|---|---|
| *Cebuella pygmaea* | KC757389 | NA | NA | NA | NA | NA |
| *Cebus albifrons* | NC_002763 | NA | NA | NA | NA | NA |
| *Chiropotes albinasus* | KC757393 | NA | NA | NA | NA | NA |
| *Lagothrix lagotricha* | KC757398 | NA | NA | NA | NA | NA |
| *Leontopithecus rosalia* | KC757399 | NA | NA | NA | NA | NA |
| *Plecturocebus cupreus* | KC959986 | NA | NA | NA | NA | NA |
| *Plecturocebus donacophilus* | FJ785423 | NA | NA | NA | NA | NA |
| *Saguinus oedipus* | KC757409 | NA | NA | NA | NA | NA |
| *Saimiri boliviensis boliviensis* | NC_018096 | NA | NA | NA | NA | NA |
| *Saimiri oerstedii citrinellus* | HQ644336 | NA | NA | NA | NA | NA |
| *Saimiri sciureus macrodon* | HQ644338 | NA | NA | NA | NA | NA |
| *Saimiri oerstedii oerstedii* | HQ644337 | NA | NA | NA | NA | NA |
| *Saimiri boliviensis peruviensis* | HQ644340 | NA | NA | NA | NA | NA |
| *Saimiri sciureus sciureus* | HQ644334 | NA | NA | NA | NA | NA |
| *Sapajus apella* | NC_016666 | NA | NA | NA | NA | NA |
| *Sapajus xanthosternos* | KC757410 | NA | NA | NA | NA | NA |

Table S2. Each cell lists individuals that possess the same mtDNA haplotypes.

| |
|---|
| BJT6, BJ65 |
| Cge006, Cgoeffroyi NC21941 |
| BJT107, BJT109 |
| BJT169, BJT170, BJT171 |
| BJT13, BJT14, BJT15, BJT16, BJT31, BJT38, BJT39, BJT75, BJT150, BJT151 |
| Cjacchus NC_025586, Cjacchus KM588314A |
| BJT166, BJT167 |
| BJT8, BJT10 |
| Ckuhlii NC_027658, Ckuhlii KR869628 |

Table S3. Organization of the *C. aurita* mitochondrial genome based on 16,471 sequenced bases of individual BJT065 (Accession number MT041703).

| Region | Position | | Length | Strand | Codon | |
|---|---|---|---|---|---|---|
| | From | To | | | Start | Stop |
| tRNA-Phe | 1 | 69 | 69 | H | | |
| 12s-rRNA | 70 | 1019 | 950 | H | | |
| tRNA-Val | 1024 | 1091 | 68 | H | | |
| 16s-rRNA | 1092 | 2644 | 1553 | H | | |
| tRNA-Leu (UUR) | 2646 | 2720 | 75 | H | | |
| ND1 | 2723 | 3679 | 957 | H | GTG | TAA |
| tRNA-Ile | 3679 | 3747 | 69 | H | | |
| tRNA-Gln | 3745 | 3816 | 72 | L | | |
| tRNA-Met | 3820 | 3887 | 68 | H | | |
| ND2 | 3890 | 4928 | 1039 | H | ATC | TAG |
| tRNA-Trp | 4929 | 4995 | 67 | H | | |
| tRNA-Ala | 5004 | 5072 | 69 | L | | |
| tRNA-Asn | 5074 | 5146 | 73 | L | | |
| tRNA-Cys | 5179 | 5245 | 67 | L | | |
| tRNA-Tyr | 5245 | 5311 | 67 | L | | |
| COI | 5319 | 6875 | 1557 | H | ATG | AGG |
| tRNA-Ser(UCN) | 6864 | 6931 | 68 | L | | |
| tRNA-Asp | 6936 | 7003 | 68 | H | | |
| COII | 7004 | 7691 | 688 | H | ATG | TA |
| tRNA-Lys | 7692 | 7758 | 67 | H | | |
| ATP8 | 7759 | 7965 | 207 | H | GTG | TAG |
| ATP6 | 7920 | 8600 | 681 | H | ATG | TAA |
| COIII | 8600 | 9383 | 784 | H | ATG | T |
| tRNA-Gly | 9384 | 9450 | 67 | H | | |
| ND3 | 9451 | 9796 | 346 | H | ATA | T |
| tRNA-Arg | 9797 | 9862 | 66 | H | | |
| ND4L | 9866 | 10162 | 297 | H | ATG | TAA |
| ND4 | 10156 | 11530 | 1375 | H | ATG | T |
| tRNA-His | 11531 | 11599 | 69 | H | | |
| tRNA-Ser(AGY) | 11600 | 11658 | 59 | H | | |
| tRNA-Leu(CUN) | 11659 | 11729 | 71 | H | | |

| | | | | | | |
|---|---|---|---|---|---|---|
| ND5 | 11733 | 13538 | 1806 | H | ATA | TAA/TAG |
| ND6 | 13535 | 14068 | 534 | L | ATG | TAA |
| tRNA-Glu | 14069 | 14137 | 69 | L | | |
| CYT B | 14142 | 15281 | 1140 | H | ATG | TAG |
| tRNA-Thr | 15284 | 15352 | 69 | H | | |
| tRNA-Pro | 15354 | 15422 | 69 | L | | |
| Control region | 15427 | 16471 | 1045 | H | | |

Table S4. Divergence times for *Callithrix* species and select nodes (MRCA = Most recent common ancestor; values in brackets = 95% highest posterior density). Node names follow major node designations shown in Figure S3 as capital letters.

| Node | Taxa Diverging at Node | Divergence Time |
|---|---|---|
| A | *C. jacchus* - *C. penicillata* (Caatinga clade) | 0.51 [0.35-0.69] |
| B | *C. kuhlii* - (*C. jacchus* + *C. penicillata* (Caatinga clade)) | 0.82 [0.59-1.09] |
| C | *C. geoffroyi* - (*C. kuhlii* + *C. jacchus* + *C. penicillata* (all clades)) | 1.18 [0.87-1.58] |
| D | *C. aurita* - (*C. geoffroyi* + *C. kuhlii* + *C. jacchus* + *C. penicillata* (all clades)) | 3.54 [2.37-4.88] |
| E | *Callithrix* - (*Cebuella/Mico*) | 6.83 [4.86-9.39] |
| F | *Saguinus*- (*Cebuella*+ *Callithrix*) | 13.42 [9.66,18.82] |
| G | *Leontopithecus* -(*Saguinus* + *Cebuella*+ *Callithrix*) | 15.69 [11.73-22.06] |
| H | Callitrichidae -(*Aotus*+*Sapajus*+*Saimiri*) | 18.63 [15.28-22.80] |
| I | Atelidae - (Cebidae + Callitrichidae) | 19.76 [16.43-23.48] |
| J | Pithecidae - (Atelidae + Cebidae + Callitrichidae) | 21.68 [18.56-27.23] |

Table S5. BMM posterior probabilities for Figure 4 nodes. Location abbreviations follow Figure 4.

| Node | Location(s) | PP | Location(s) | PP | Location(s) | PP | Location(s) | PP | Location(s) | PP |
|---|---|---|---|---|---|---|---|---|---|---|
| 60 | AF | 99.93% | A | 0.03% | F | 0.02% | | | | |
| 61 | AF | 99.92% | A | 0.03% | F | 0.02% | | | | |
| 62 | AF | 99.89% | A | 0.04% | F | 0.02% | | | | |
| 63 | AF | 99.55% | A | 0.17% | F | 0.16% | | | | |
| 64 | AF | 98.95% | F | 0.41% | A | 0.20% | | | | |
| 65 | AF | 54.17% | F | 41.51% | A | 2.20% | | | | |
| 66 | F | 49.68% | AF | 44.83% | A | 3.82% | | | | |
| 67 | F | 76.88% | AF | 19.73% | A | 1.98% | | | | |
| 68 | AF | 44.21% | F | 34.13% | A | 18.45% | | | | |
| 69 | G | 99.56% | FG | 0.08% | CG | 0.05% | | | | |
| 70 | G | 99.20% | FG | 0.18% | DG | 0.07% | | | | |
| 71 | G | 99.59% | FG | 0.08% | CG | 0.05% | | | | |
| 72 | G | 99.21% | FG | 0.19% | BG | 0.07% | | | | |
| 73 | G | 96.14% | FG | 1.28% | F | 0.37% | | | | |
| 74 | G | 35.60% | F | 20.60% | A | 15.14% | H | 5.64% | | |
| 75 | H | 97.77% | GH | 0.66% | CH | 0.18% | | | | |
| 76 | H | 50.26% | G | 11.68% | P | 6.11% | | | | |
| 77 | P | 98.16% | OP | 0.47% | DP | 0.16% | | | | |
| 78 | P | 48.67% | O | 13.39% | H | 8.92% | | | | |
| 79 | O | 99.62% | DO | 0.05% | EO | 0.05% | | | | |
| 80 | O | 99.41% | C | O 0.06% | D | O 0.06% | | | | |
| 81 | O | 98.13% | C | O 0.21% | B | O 0.21% | | | | |
| 82 | O | 50.03% | P | 7.53% | I | 7.13% | J | 6.75% | | |
| 83 | IJK | 92.26% | IJ | 7.14% | JK | 0.16% | | | | |
| 84 | IJ | 32.41% | IJK | 30.46% | J | 16.76% | JK | 15.75% | | |
| 85 | I | 91.93% | I | J 4.03% | IK | 3.13% | | | | |
| 86 | IJ | 32.14% | IJK | 31.57% | I | 16.25% | IK | 15.96% | | |
| 87 | IJ | 42.25% | IJK | 34.84% | I | 6.76% | IK | 5.57% | | |
| 88 | I | 31.89% | J | 29.56% | K | 10.09% | | | | |
| 89 | L | 60.82% | M | 14.23% | N | 10.71% | | | | |
| 90 | L | 44.43% | M | 38.14% | LM | 10.73% | | | | |
| 91 | L | 64.02% | M | 21.05% | LM | 9.09% | | | | |
| 92 | M | 97.77% | LM | 1.09% | L | 0.21% | | | | |
| 93 | M | 70.13% | L | 15.29% | LM | 4.62% | | | | |
| 94 | M | 26.09% | I | 12.39% | J | 11.70% | L | 8.59% | K | 6.56% |

Table S6. Summary of record numbers for collected samples that have been entered into the Brazilian CGEN SISGEN sample database (ES = Espírito Santo, MG = Minas Gerais, PE = Pernambuco, RJ = Rio de Janeiro, SP = São Paulo).

| City | State | Wild Animal | Translocated/ Captive | Facility | SISGEN Record Number |
|---|---|---|---|---|---|
| Santa Teresa | ES | x | | | ABD1A11 |
| Serra | ES | x | | | AE61F27 |
| Guiricema | MG | x | | | A94942B |
| Berilo | MG | x | | | A3CB657 |
| Lavras | MG | x | | | A679090 |
| Belo Horizonte | MG | x | | | A679090 |
| Vicosa | MG | x | | | A998C84 |
| Petrolina | PE | | x | CEMAFAUNA | A3F0F46, A679090 |
| Guapirim | RJ | | x | Rio de Janeiro Primate Center | A2AEE0E, AD432CF, A6F3E89, A3F0F46, A6F3E89 |
| Campos | RJ | | x | SERCAS | AD2F796 |
| Silva Jardim | RJ | x | | | A4A29D7 |
| Guarulhos | SP | | x | Guarulhos Zoo | A18C1CE, A3F0F46, A679090, A20E733 |

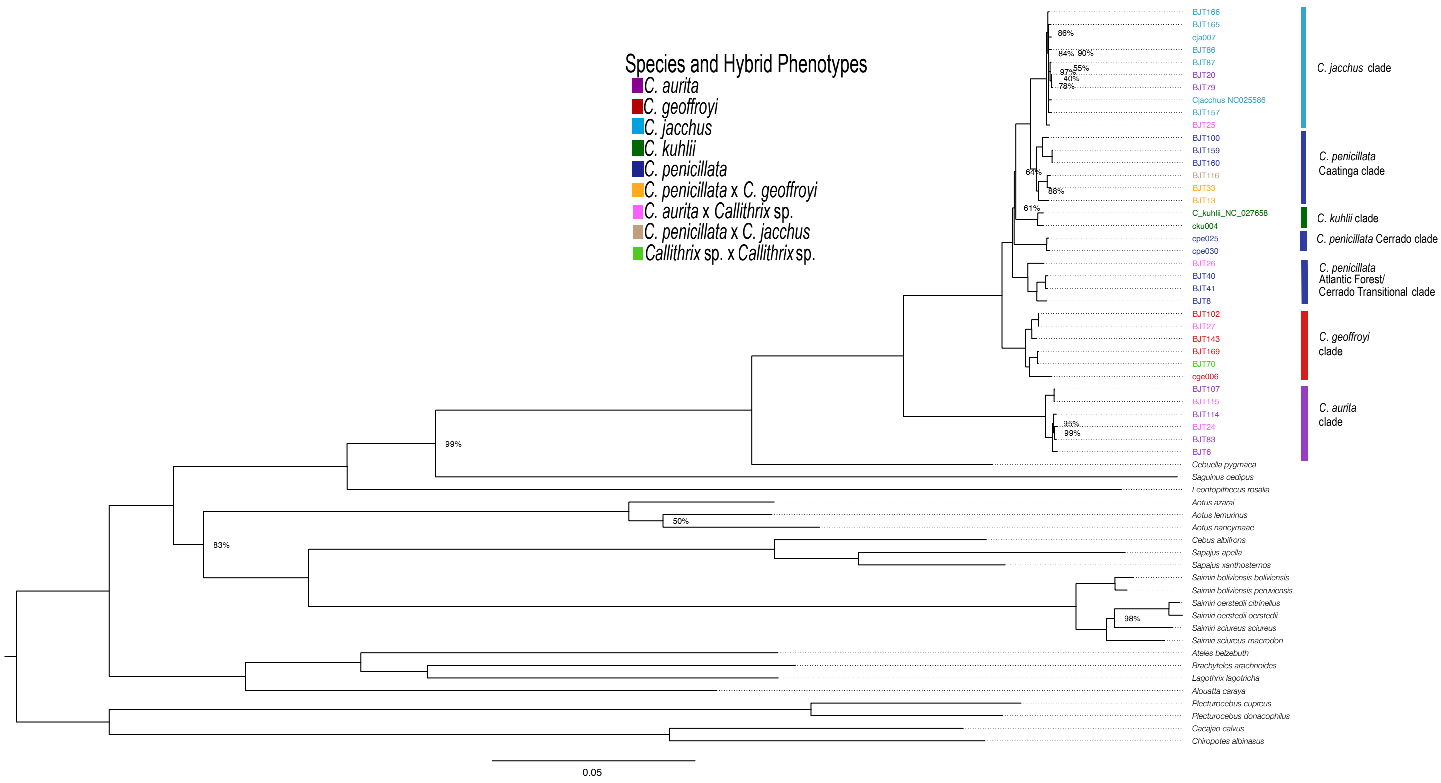


Figure S1. Maximum-likelihood (ML) tree showing phylogenetic relationships among *Callithrix* haplotypes as calculated from mtDNA genome sequences. Numbers at nodes indicate bootstrap support for a given node, otherwise node bootstrap support was 100%. Haplotype colors at tips correspond to the 'Species and Hybrid Phenotypes' legend, and indicate phenotypes associated with each given haplotype.

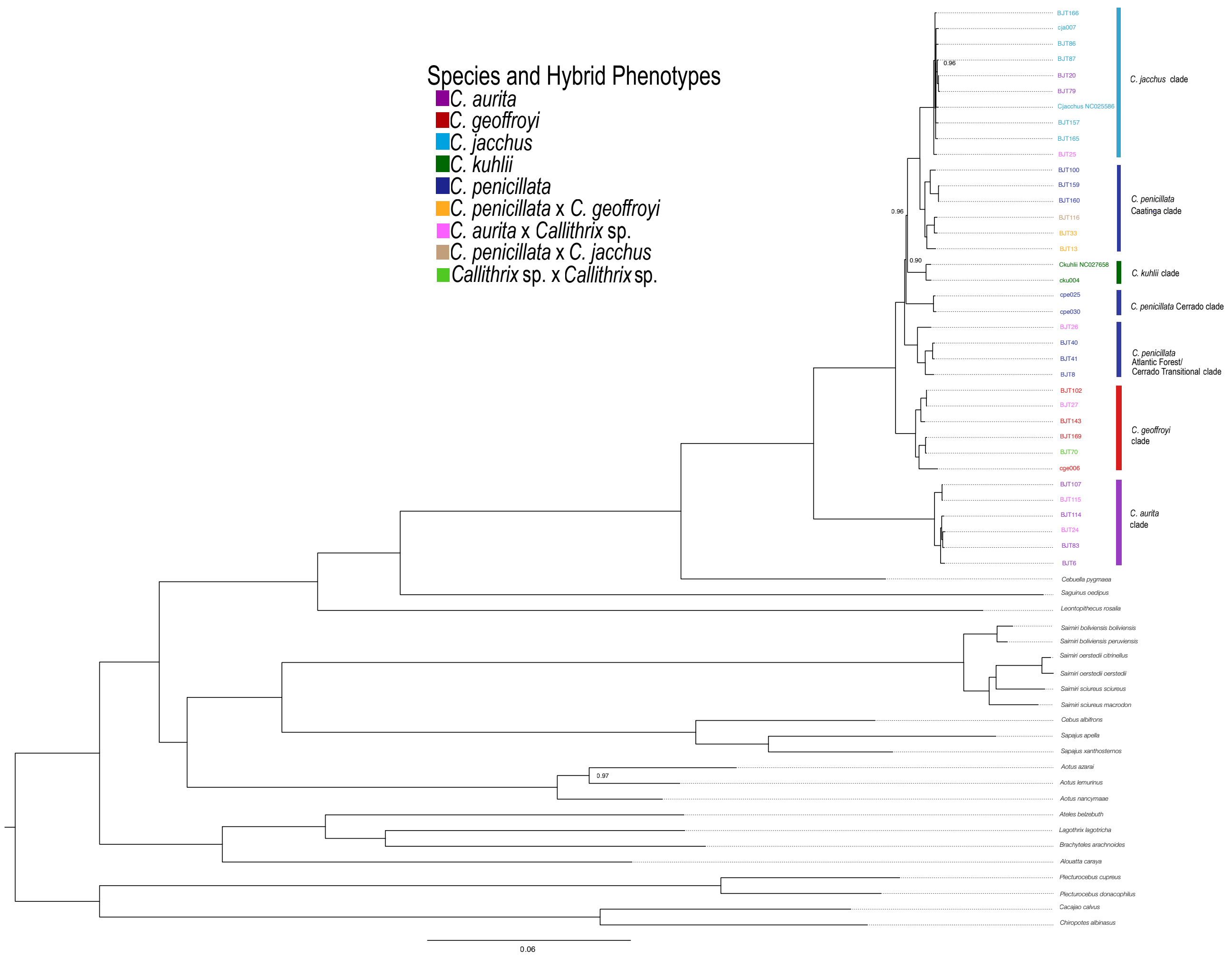


Figure S2. Bayesian tree showing phylogenetic relationships among *Callithrix* species and hybrid haplotypes from mtDNA genome sequences. Numbers at nodes indicate posterior probability for a given node, otherwise node posterior probability was 1. Haplotype colors at tips correspond to the 'Species and Hybrid Phenotypes' legend, and indicate phenotypes associated with each given haplotype.

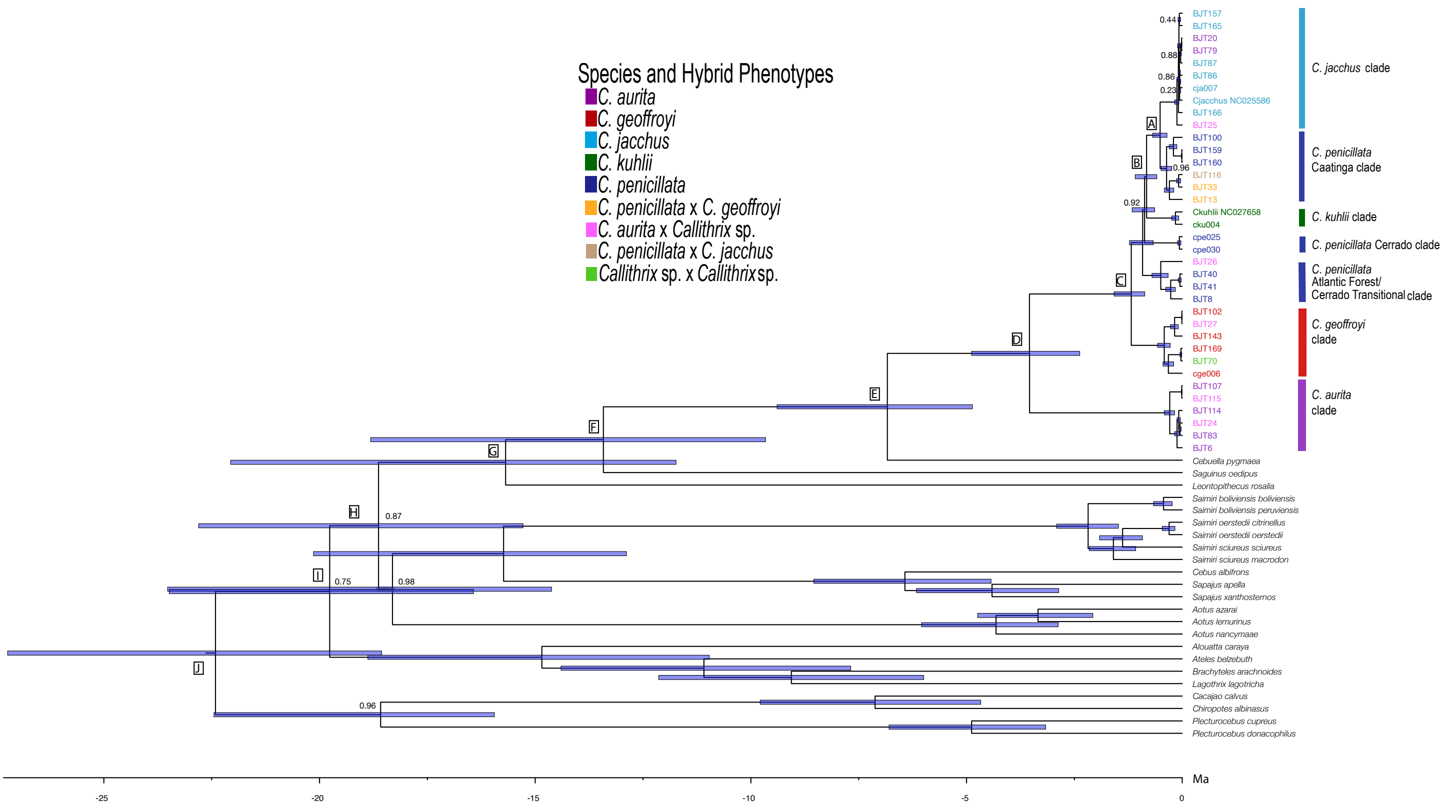


Figure S3. BEAST tree showing phylogenetic relationships and divergence ages in million years (Ma) among *Callithrix* haplotypes and other New World primates as calculated from mtDNA genome sequences. Major nodes are identified by capital letters, and blue bars at all nodes indicate 95% highest posterior densities (HPD) of divergence times. Haplotype colors at tips correspond to the 'Species and Hybrid Phenotypes' legend, and indicate phenotypes associated with each given haplotype.
